# Membrane potential mediates the cellular response to mechanical pressure

**DOI:** 10.1101/2023.11.02.565386

**Authors:** Avik Mukherjee, Yanqing Huang, Jens Elgeti, Seungeun Oh, Jose G. Abreu, Anjali Rebecca Neliat, Janik Schüttler, Dan-Dan Su, Christophe Dupre, Nina Catherine Benites, Xili Liu, Leonid Peshkin, Mihail Barboiu, Hugo Stocker, Marc W. Kirschner, Markus Basan

## Abstract

Mechanical forces have been shown to influence cellular decisions to grow, die, or differentiate, through largely mysterious mechanisms. Separately, changes in resting membrane potential have been observed in development, differentiation, regeneration, and cancer. We now demonstrate that membrane potential is the central mediator of cellular response to mechanical pressure. We show that mechanical forces acting on the cell change cellular biomass density, which in turn alters membrane potential. Membrane potential then regulates cell number density in epithelia by controlling cell growth, proliferation, and cell elimination. Mechanistically, we show that changes in membrane potential control signaling through the Hippo and MAPK pathways, and potentially other signaling pathways that originate at the cell membrane. While many molecular interactions are known to affect Hippo signaling, the upstream signal that activates the canonical Hippo pathway at the membrane has previously been elusive. Our results establish membrane potential as a central regulator of growth and tissue homeostasis.

## INTRODUCTION

Cells in confluent tissues need to precisely control biomass synthesis and proliferation to quickly heal injuries^1–3^, while avoiding the excessive growth that is the hallmark of tumorigenesis^4,5^. Mechanical signaling has been proposed to be involved in this process^6–17^ and mechano-sensitive signaling pathways like the Hippo pathway have been shown to have important roles^18–22^. However, the connection between mechanical signaling and tissue density homeostasis is not clear, even in principle. Accurate cellular decision-making about growth or survival requires information on physiological parameters on the cellular and tissue scale, such as cell volume or total cellular biomass, both in relation to the available space in the tissue. Local mechanisms of mechanotransduction do not provide satisfying explanations for homeostatic control of cell size and growth. For example, tension in the plasma membrane results in calcium influx via stretch-activated channels^23–25^. However, providing information for cell and tissue growth on long timescales, would require precise control of the total amount of plasma membrane in relation to cellular biomass as cells change in size and shape. To the contrary, mechanical membrane stretching readily results in production of additional membrane, showing that the amount of plasma membrane is not constrained in this way^26,27^. Similarly, tension on individual cell-cell adherence junctions and cytoskeletal bonds can mediate mechanical signaling. However, these forces depend on cell shape and cytoskeletal contractility and it is unclear if they can be used as a readout of total cell size and cellular biomass. We therefore set out to identify additional cellular variables that could connect tissue-level spatial constraints to cellular growth control.

Cellular biomass density is generally assumed to be constant for a given cell type, implying that cell mass and volume are directly proportional, even as cells undergo additional rounds of proliferation in already confluent epithelia^28^ or when cell volume changes quickly and dramatically during wound healing. By directly measuring cellular biomass density, we discovered that this assumption is incorrect. Instead, we find that biomass density is a central physiological variable that is intimately coupled to cellular growth control and transduction of mechanical pressure. We reveal a direct one-to-one coupling from mechanical forces acting on cells, to changes in cytoplasmic biomass density, to cellular resting membrane potential that controls signal transduction. The mechanism of this coupling is distinct from and orthogonal to known mechanisms of mechano-transduction like calcium signaling or tension on adherence junctions. Instead, it emerges from the intrinsic mechano-electro-osmotic properties of the cytoplasm and from the coupling of biomass counterions with membrane potential. This mechanism provides cells with a global, instantaneous readout of spatial constraints in tissues in relation to their biomass and mechanical forces acting on them. This readout is independent of cell size and shape and invariant under changes that strongly affect other mechanisms of mechanotransduction, such as surface-to-volume ratio and the amount of plasma membrane of the cell. To our knowledge, this is the first example of a self-contained transduction mechanism for mechanical pressure that enables cells to determine their cytoplasmic biomass density. We show that this mechanism plays a central role in controlling growth and homeostasis in confluent tissues.

## RESULTS

### Cellular biomass density increases with tissue density

To measure cellular biomass density, we used normalized Raman imaging (NoRI)^29^, which enables absolute quantification of cellular macromolecular components and water. Specifically, we aimed to quantify intracellular biomass densities at increasing levels of confluence. We used different cell lines that model epithelial monolayers^30^ of varying tissue origin, including MDCK, MCF10a, and EpH4-Ev cells. In all cases, we observed that cellular biomass density underwent a striking, previously unobserved, increase as a function of cell number density. Fig. 1a shows the nuclear density and biomass density of MDCK cells at 0 hours, 48 hours, and 96 hours after initially reaching confluence, and Fig. S1a-d shows the separate protein and lipid contributions. Fig. S2a-b shows increase in cellular biomass density as a function of time after reaching confluence. We observed a similar trend in MCF10a and EpH4-Ev monolayers (Fig. 1c, 1d, Fig. S1e, S1f). We confirmed this observation with an independent microscopic technique based on laser interferometry, referred to as holo-tomographic microscopy (Tomocube) (Fig. 1a, red squares & Fig. S2 e-f)^31,32^.

**Figure 1:**
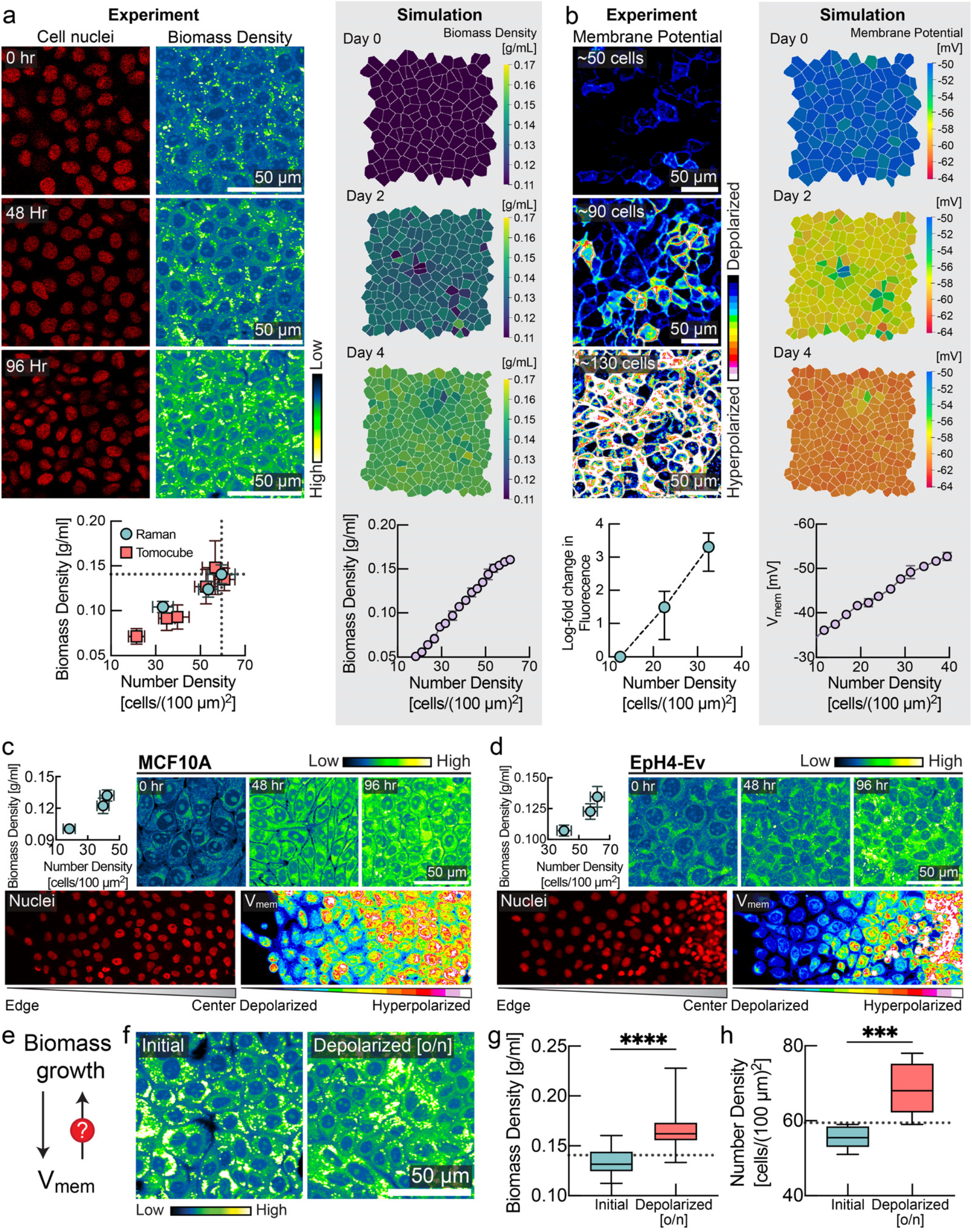
Membrane potential is a sensor and regulator of tissue density. **a**, Biomass density increases with cell density. MDCK cells were imaged for 4 consecutive days after confluence with increasing cell number density. Left, number of cell nuclei and cellular biomass density (in units of g/ml) measured via Normalized Stimulated Raman Spectroscopy (NoRI). Below, correlation between number density (mean ± s.d., NoRI: n=8, Tomocube: n=10) and biomass density (mean ± s.d., NoRI: n=15, Tomocube: n=∼20) measured using NoRI and Tomocube. Right, snapshots of biomass density in tissue simulation (mean ± s.d., n=4 simulations). **b**, Membrane hyperpolarization increases with cell density. Left, fluorescence of MDCK-Voltron cell. Below, log fold change in fluorescence (mean ± s.d., n=8) against number density. Right, snapshots of membrane potential in tissue simulation (mean ± s.e.m., n=4 simulations). **c-d**, Correlation between cell density and membrane potential in (c) MCF10A and (d) EPH4-Ev cells. Top panels: NoRI images and quantification (mean ± s.d., number density: n=∼10, NoRI: n=24) of MCF10A/EpH4-Ev cells for 4 consecutive days after confluence with increasing cell number density. Bottom panels: DiOC_2_(3) imaging from colony edge to colony centre. **e**. Feedback control loop between biomass growth and membrane potential is crucial for epithelial homeostasis. **f-h**, Depolarization of membrane potential by ouabain increases the biomass density (box plot, line at mean, error bars represent min to max, n=40) and number density (n = 8) of confluent cells over the homeostatic steady state limit (dotted line represents the mean homeostatic cell number density and biomass density; quantified from experiment in panel a).

### Cell membrane potential increases with increasing tissue density

Cellular biomass components, including proteins, RNA and DNA, carry a net negative electric charge at physiological pH^33–35^. Inspired by the Gibbs-Donnan effect, a phenomenon from equilibrium thermodynamics in which a membrane that is permeable to some charged species and not others can create an asymmetric distribution of charged particles^36^, we asked whether changes in biomass density were correlated with changes of membrane potential. Using the hybrid voltage-indicator Voltron^37^, we quantified membrane potential in MDCK cells as a function of cell number and biomass density. Strikingly, MDCK cells became increasingly hyperpolarized with increasing cell number (Fig. 1b). In MCF10a and EpH4-Ev cells, we also confirmed the hyperpolarization of membrane potential with increasing cell number density (Fig. 1c and Fig. 1d, lower panel) using the membrane potential dye DiOC_2_(3). These findings are consistent with observations from the 1970s of a correlation between cell density and membrane potential^38^. This earlier study used glass-microelectrode-based measurements of resting membrane potential in CHO cells as they reached different levels of confluence over time and in different regions of a colony with varying local densities of cells. The mechanistic basis and physiological consequences of these early observations has never been determined. We recapitulated these measurements and find that these changes in membrane potential (Fig. S3a,b,d) directly correspond to changes in cellular biomass density (Fig. S3c,e). While MDCK and MCF10a cells form a tight epithelium connected by tight junctions, CHO cells do not, suggesting that the relationship between membrane potential and biomass density is at least partly cell-autonomous.

The correlation between biomass density and membrane potential indicated to us that these physiological variables might be directly coupled. To test this, we applied external osmotic pressure to compress cells, artificially increasing their biomass density. Indeed, we found that the membrane potential of CHO and MDCK cells hyperpolarized with mechanical compression (Fig. S4). Therefore, changes in membrane potential could be a direct result of changes in biomass density.

### Membrane potential regulates biomass density and tissue density

Based on these observations, we hypothesized that membrane potential could be directly involved in regulating biomass production and proliferation (Fig. 1e). To test this hypothesis, we first grew MDCK monolayers to their maximum density (Fig. 1f). We confirmed that cells reached homeostatic biomass density using NoRI (Fig. 1f, 1g). Next, we depolarized the cells overnight by inhibiting the sodium-potassium pump using the drug ouabain. Indeed, we found that depolarization pushed cells to a significantly higher biomass density relative to their normal homeostatic biomass density (Fig 1f,g) and significantly higher cell numbers (Fig. 1h). We confirmed that this phenotype was independent of the pharmacological method of depolarization in further experiments. We grew MDCK cells to maximum density and then applied different depolarizing agents (gramicidin, or the synthetic gramicidin analog TCT^39,40^) or hyperpolarizing drugs (valinomycin) at different concentrations to tissue culture. Cells in depolarizing conditions (Fig. S5a, red) maintained a significantly higher cell number density than cells grown in the absence of drugs (Fig. S5a, cyan). Conversely, the cell number density in hyperpolarizing conditions (Fig. S5a, yellow) dropped substantially and then plateaued at a much lower level. We found that steady-state cell number density could be continuously modulated by varying the dose of the hyperpolarizing drug (Fig. S5c).

### Mechano-electro-osmotic model

To rationalize the effect of biomass density on membrane potential, we formulated a biophysical mathematical model of ion homeostasis, cellular biomass and mechanical pressure based on first principles (Box 1). Classical models of membrane potential like the Goldman-Hodgkin-Katz model do not consider mechanical forces at all, and their systems of equations are only closed by introducing convenient, yet unphysical simplifications^41,42^. Models more correctly rooted in elementary physical principles that consider active ion transport and ion leakage were developed in the 1960s and showed that osmotic stabilization of cells requires constant ion export that results in a build-up of membrane potential as a by-product^43–45^. However, these models did not consider growing cells, which change in cell mass, cell membrane, cell volume and cell shape, affecting model parameters like permeability and transport activity, and therefore did not address the connection between membrane potential and growth control. It has also been generally assumed that cellular biomass density^28^ and membrane potential^46^ are tightly regulated and constant for a given cell type. Thus, the central hypothesis of our work, that the elementary biophysical coupling between biomass density and membrane potential can constitute a mechanism of signal transduction, has never been explored.

Membrane potential is intimately coupled to the balance of charged ions and macromolecules inside the cell, which also controls the osmotic pressure of the cytoplasm. As illustrated in Box 1 (top), a cell resists external forces compressing it via osmotic pressure. This results from increased concentrations both of the intracellular macromolecules in the cytoplasm and of their counterions. A substantial fraction of these counterions are not condensed on macromolecules and are therefore osmotically active even at very high salt concentrations, as demonstrated by classical experimental and theoretical work^47–49^. Of note, the net charge on the macromolecules in the cell is negative because the proteome has an average isolectric point above 7.0 and nucleic acids are negatively charged. In particular, the significant fraction of the biomass in the cytoplasm comprised of RNA^50^ is negatively charged. Thus, external pressure results in a more concentrated cytoplasm and a steeper concentration gradient across the plasma membrane, promoting a larger efflux of positively charged ions by diffusion. A buildup of membrane potential due to the retention of negatively charged biomolecules inside the cell counteracts this diffusive efflux. In equilibrium conditions, this is known as the Gibbs-Donnan effect^36^, but the same processes also function far from equilibrium.

Overall, the model shows that when the expression ratio of ion transporters and channels is constant (e.g. because pumps and channels are expressed at a constant fraction of the proteome, which is arguably the simplest possible assumption), membrane potential constitutes a cell-size-independent readout of total mechanical pressure on the cell and the corresponding cytoplasmic biomass density (see Box 1 & Supplementary Note 1 for details of the mathematical model). No specific scaling of expression levels with membrane area, cell volume, cell mass or time is required. In effect, membrane potential physically controls biomass density and mechanical forces, and vice versa. The consequence is that membrane potential offers a near-instant integrated readout of biomass density and mechanical forces that is unperturbed by growth, cell size and cell shape (Fig. 1a-b).

### Tissue simulation

We calibrated the mechano-electro-osmotic model with physiological parameters and used it to simulate a mono-layered epithelium (Supplementary Note 2). To test the idea that tissues can achieve homeostasis relying only on the membrane potential as a control variable, we developed a model where the membrane potential is the sole regulator of biosynthesis and apoptosis (see Supplementary Note 2 for details of implementation).

This tissue simulation successfully recapitulated our experimental observations, including the increase in biomass density with cell number density (Fig. 1a, right panel, and Fig. S2 c-d) and the corresponding changes in membrane potential (Fig. 1b, right panel). Implementing the mechanism of different depolarizing and hyperpolarizing drugs in the simulation, the model recapitulated the observed changes in steady-state tissue density (Fig. S5b). The general effect of these drugs can be understood intuitively: cells perceive mild depolarization to be a state of low confluence. They therefore continue to grow, generating an abnormally high density (Fig. S5b, red circles). This is remarkable and non-trivial, given that depolarization should normally result in swelling and thereby a decrease in cellular biomass density. Conversely, hyperpolarization caused by drugs leads to a lower homeostatic tissue density because cells interpret hyperpolarization as excess tissue density and arrest their growth prematurely (Fig. S5b, yellow circles). Thus, our model of regulation of cell growth by membrane potential correctly predicts changes in steady-state tissue density.

**Box 1:**
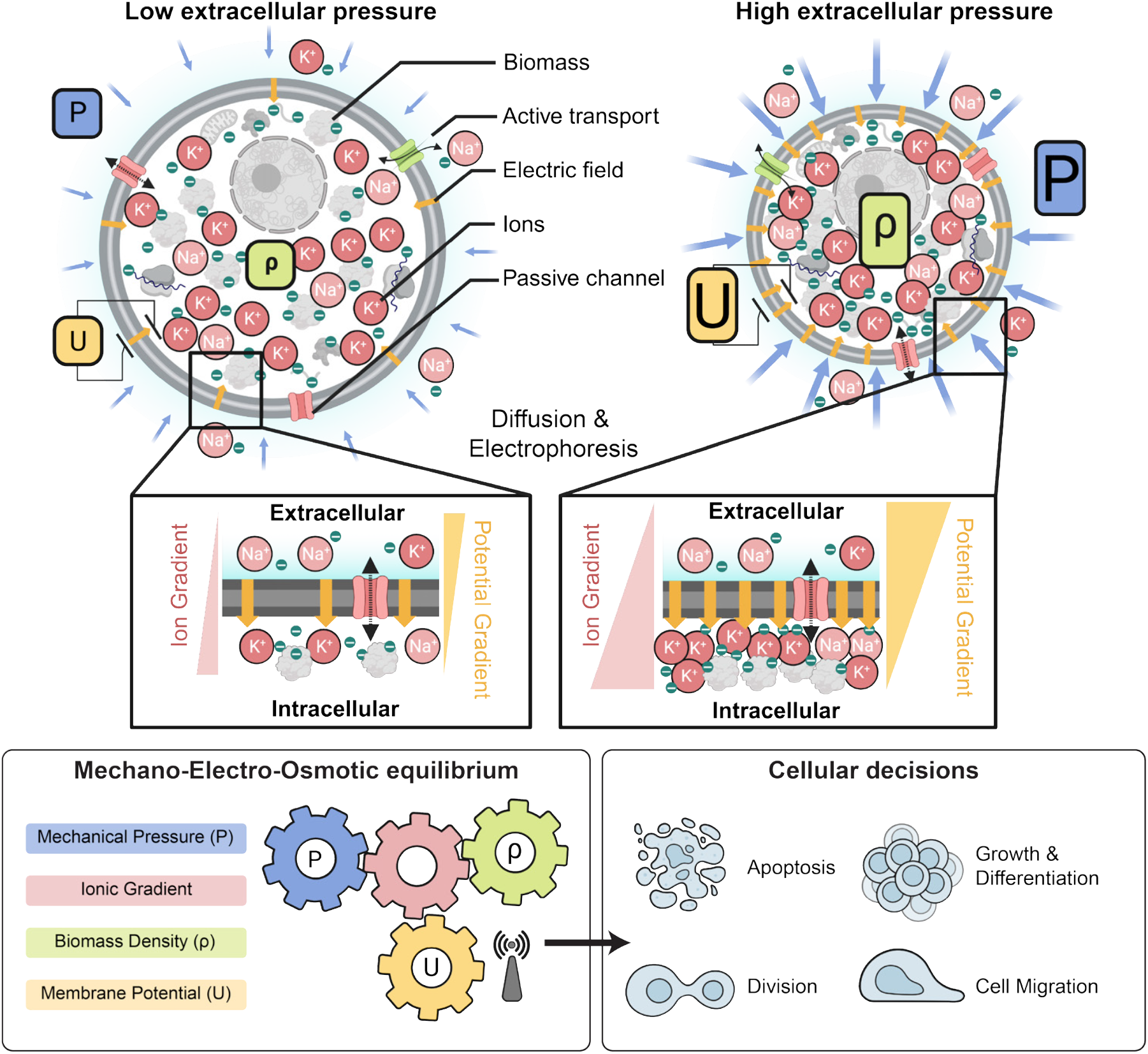
Mechano-electro-osmotic model. Top: The model combines three elementary conservation laws: Flux balance across the plasma membrane, mechanical force balance, and charge balance. The left-hand side illustrates a cell under low mechanical pressure and the right-hand side a corresponding cell under high mechanical pressure. Cellular biomass like protein and RNA carries a net negative charge. Cells resist compression of their volume via cytoplasmic osmotic pressure that counteracts and balances external forces (force balance). This osmotic pressure originates from an intracellular concentration of counterions that are attracted to balance negatively charged macromolecules (charge balance). As illustrated on the right-hand side, higher mechanical pressure acting on the cell is thus balanced by a higher concentration of counterions. Higher intracellular ion concentrations result in steeper ion concentration gradients across the membrane, as illustrated in the right inset. Thus, with constant ion channel abundances, a steeper concentration gradient across the membrane results in an augmented diffusive efflux of ions (Fick’s law). With constant active ion transport flux (sodium-potassium-ATPase), this increased diffusive ion efflux results in a buildup of net negative charge in the cytoplasm and a corresponding membrane potential that counteracts the diffusive efflux of positively charged ions until flux balance is achieved (flux balance for each ion species). **Bottom:** A one-to-one correspondence between mechanical pressure, cytoplasmic biomass density and membrane potential emerges from the mechano-electro-osmotic equilibrium illustrated in the top panel.

### Membrane potential reflects mechanical forces

Our mechano-electro-osmotic model (Box 1) predicts that mechanical forces should be directly reflected in the membrane potential. To test this prediction, we imposed external mechanical perturbations on the tissue and measured the response of membrane potential, using an elastic membrane that can be stretched or de-stretched mechanically, thereby expanding or compressing cells growing on the membrane^51,52^ (Fig. 2a). We cultured MDCK cells on a stretchable PDMS membrane and waited until cells reached maximum cell number density. We used the voltage sensitive dye FluoVolt, whose fluorescence intensity increases with depolarization^53^, to image changes in membrane potential on short timescales. Stretching the MDCK monolayer did result in depolarization (Fig. 2a,b, quantified in panel c), consistent with our simulation prediction (Fig. 2d). Next, we grew cells to confluence on a membrane, and applied a 20% uni-axial stretch. We again observed depolarization of the membrane potential, measured using a different voltage-sensitive dye, DiSBAC_2_(3)^54^. Reducing the stretch of the PDMS membrane quickly led to repolarization of membrane potential (Fig. 2e,f), as predicted by the simulation (Fig. 2g). Using NoRI imaging, we quantified biomass density of the confluent monolayer just before and immediately after a 20% uniaxial stretch, and then again after overnight cultivation (Fig. 2h,i). As expected from the tissue simulation (Fig. 2j), we found that stretching resulted in a significant drop in biomass density and induced biomass production and proliferation. Conversely, we also confirmed that de-stretching the PDMS membrane resulted in an increase of cellular biomass density after overnight recovery (Fig. S6a-b). These results are consistent with our hypothesis that depolarization upon stretch is caused by a drop in cellular biomass density due to an increased cell volume, rather than by tension in the plasma membrane or changes in cell-cell adhesion.

**Figure 2:**
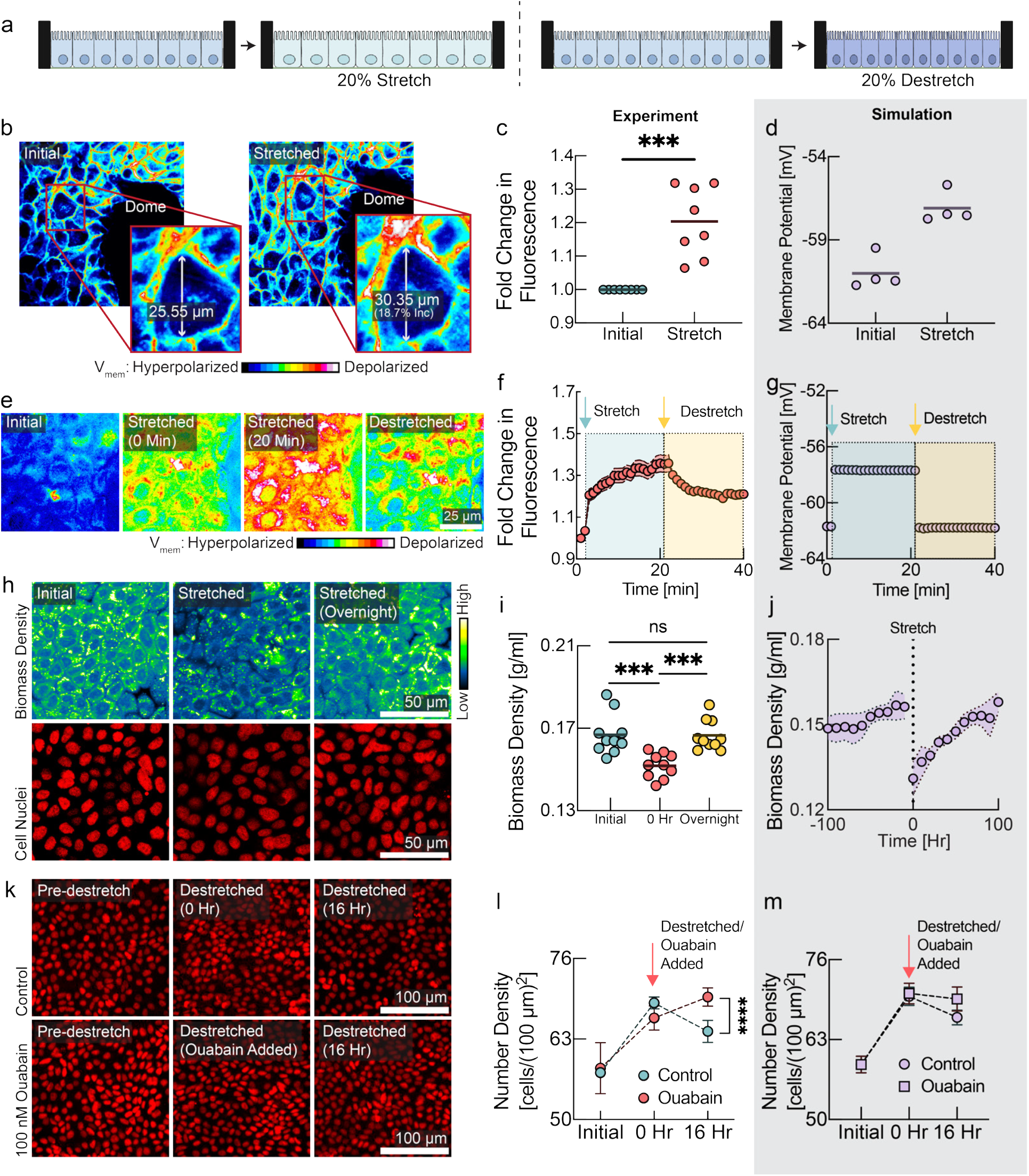
Cells detect and respond to mechanical forces via changes in membrane potential. **a**, Schematic diagram depicting the experimental design with monolayers grown on stretchable membrane. Left: cells were grown to confluence and then 20% uniaxial stretch was applied. Right: cells were grown to confluence on a pre-stretched membrane, then the stretch was released to induce compression. **b**, FluoVolt images of MDCK cells before and after stretching. **c**, Quantification of panel b (line at mean, n=8, p=0.0006, paired t-test). **d**, Membrane potential predicted by the tissue simulation upon a 20% uniaxial stretch (line at mean, n=4 simulations, each point is averaged across all cells in simulation). **e**, DiSBAC_2_(3) imaging of MDCK cells before stretching, during stretching, and after destretching showing the reversibility of membrane potential change. **f**, Quantification of panel e (mean ± s.d., n = 8). Increase in fluorescence intensity indicates depolarization and decrease indicates hyperpolarization. Time of stretch and destretch are indicated by cyan and yellow arrows respectively. **g**, Membrane potential predicted by the tissue simulation upon stretch and subsequent destretch (n=4 simulations, each point is averaged across all cells in simulation). **h**, NoRI images of cellular biomass in response to tissue stretch. Biomass density recovers to the initial biomass density level overnight. **i**, Quantification of panel h. 0 hr represents time of stretch. (n=10, p-values: 0.0006 (initial vs 0 hr), 0.0001 (0 hr vs overnight), unpaired t test). **j**, Tissue simulation recapitulates the drop of biomass density upon stretch and overnight recovery. (mean ± s.d., n=4 simulations) **k**, Rescue of destretching-induced tissue crowding by ouabain. After 16 hr, a significant number of cells had been eliminated, consistent with previous studies^51,52^. **l**, Quantification of panel k (mean ± s.e.m., n = 3 experiments of 20 ROIs each, p-value < 0.0001, unpaired t-test). **m**, Tissue simulation recapitulates cell elimination after compression and rescue from elimination by depolarizing drugs (mean ± s.e.m., n=4 simulations).

### Depolarization rescues cells from elimination after mechanical compression

As a direct test of the role of membrane potential in regulating the response of cells to mechanical perturbations, we grew MDCK cells on pre-stretched membranes to high cell number density and then released the stretch to induce cell crowding and compression of the tissue (Fig. 2k). Under normal circumstances, tissue compression results in cell extrusion^51,52^. We hypothesized that this effect might be mediated by crowding-induced hyperpolarization, and that we could prevent it by depolarizing the cells. Indeed, in the presence of depolarizing drugs, cells maintained a significantly higher cell number density after the tissue was compressed (Fig. 2k,l red vs. cyan circles). The observed experimental dynamics are consistent with our model and could be recapitulated in the tissue simulation when we simulated uniaxial compression of the tissue (Fig. 2m). In a separate experiment, we induced crowding while cells were still in growth phase and observed the same increase of steady-state cell number density when cells were treated with a different depolarizing drug, with a different mechanism of action, TCT^39,40^ (Fig. S6c-d).

### Mechanical stretch results in a wave of depolarization during wound healing caused by a drop in biomass density

Wound healing is a natural situation involving substantial dynamical changes in tissue density and mechanical forces. Any tissue can experience injuries that must be detected and repaired in a controlled fashion. During wound healing, mechanisms of tissue homeostasis are operating in high gear and we hypothesized that mechanical regulation may have an important role. To investigate the role of membrane potential in wound healing, we performed scratch wound assays on confluent MDCK monolayers. We observed a wave of depolarization in the tissue, starting at the wound’s immediate edge and moving deeper into the tissue over several hours (Fig. 3a, quantified in 3b). Depolarization at wound edges has been previously observed, but not explained^56,57^. According to our model, a drop of biomass density due to mechanical stretching of the tissue could explain depolarization. Indeed, we observed an increase in cell area in the tissue indicative of mechanical stretch (Fig. 3a, middle & 3c), as cells became motile via the epithelial mesenchymal transition (Fig. S7a-c) and moved into the wound area, pulling on the cell behind them. Using NoRI, we quantified biomass density as a function of distance from the wound edge (Fig. 3d, e & Fig. S7d). Remarkably, we observed that the depolarization wave was accompanied by a concurrent drop in biomass density. The tissue simulation with the mechano-electro-osmotic model successfully recapitulates the temporal and spatial dynamics of biomass density and membrane potential during wound healing (Fig. 3a, d, lower panels).

**Figure 3:**
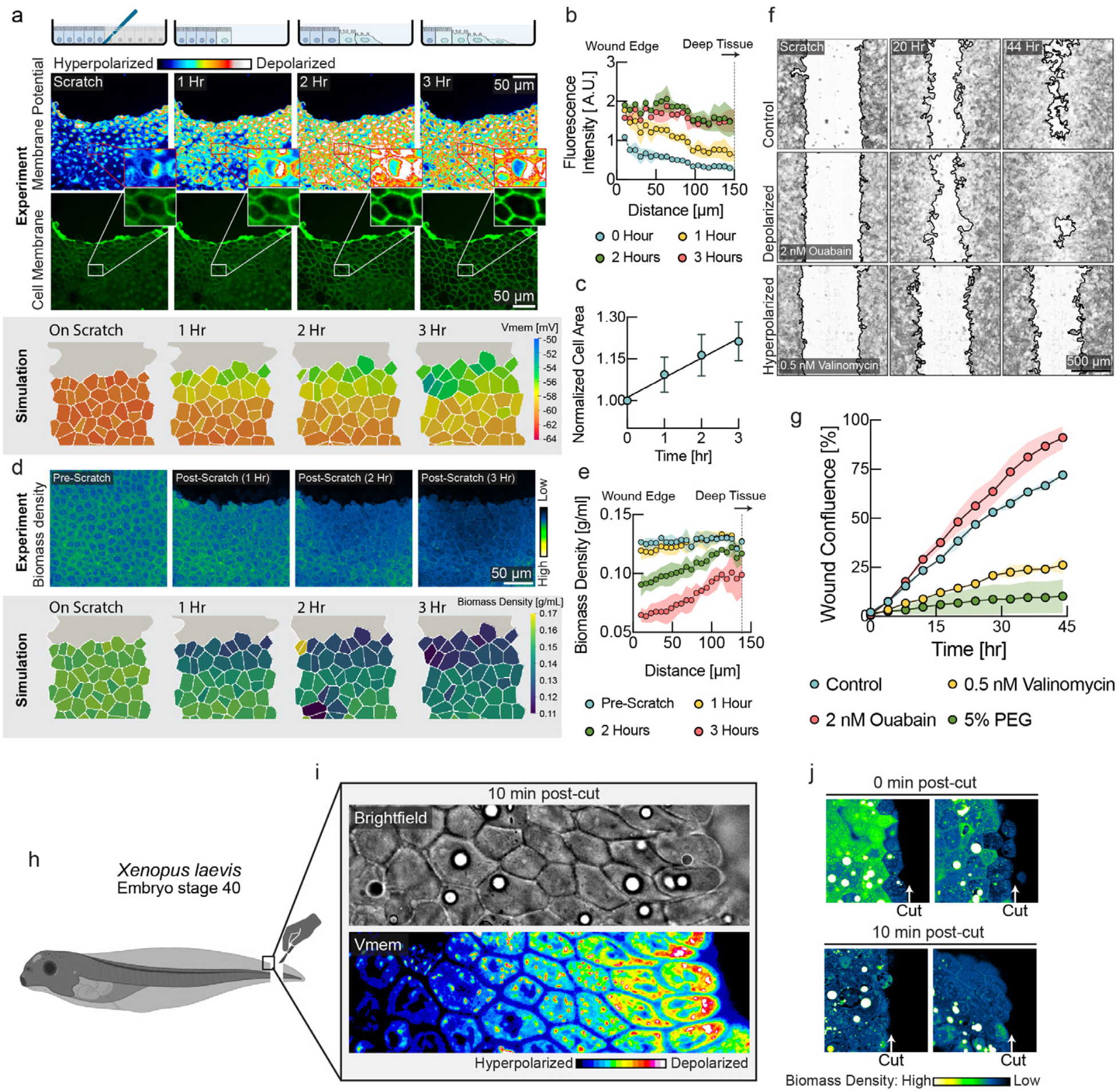
A mechanically induced depolarization wave enhances wound healing. **a**, Upper panel: DiSBAC_2_(3) and fluorescently labeled membrane images of MDCK cells imaged over 3 hours after scratch wound. A depolarization wave was observed from the wound edge up to several layers deep into the tissue. Lower panel: Membrane potential from the tissue simulation over the course of wound healing. **b**, Quantification of membrane potential as a function of distance from scratch wound at different times (mean ± s.d., n=4 rectangular ROIs of 100 µm width). **c**, Quantification of cell area shows expansion of cells during wound healing (mean ± s.d., n=9). **d**, Upper panel: NoRI images over the course of wound healing. Lower panel: Biomass density from the tissue simulation over the course of wound healing. **e**. Quantification of cellular biomass density as a function of distance from scratch wound at different times after wounding (mean ± s.d., n=4 rectangular ROIs of 50 µm width). **f**, Representative images of control, depolarizing drug treatment (ouabain) and hyperpolarizing drug treatment (valinomycin) during wound healing. Wound border outlines were generated using the Wound healing size tool^55^. **g**, Quantification of wound area closure as a function of time (mean ± s.d., n=2 wells). Depolarization resulted in faster wound healing (ouabain, red circles), as compared to the untreated control (cyan circles). Hyperpolarization (valinomycin, yellow circles, 5% PEG, green circles) resulted in slower wound healing. h, Schematic diagram showing the experimental set up for Xenopus embryo tail amputation. i, Brightfield and membrane potential (DiSBAC2(3)) images of Xenopus wound edge, 10 minutes post-amputation. Cell layers are progressively depolarized from deep tissue towards the wound edge. j, NoRI images of Xenopus tail amputation, 0 min and 10 min post-cut. The closest cell layer becomes dilute in biomass density immediately after the cut, and 5-6 cell layers become dilute in 10 minutes. Images represent different Xenopus embryos amputated and fixed at the specified times.

We hypothesized that the physiological function of this depolarization wave might be to upregulate motility and proliferation in cells that are not immediately adjacent to the wound edge. This would be expected to speed up the wound healing process. We observed both depolarization (Fig. 3a) and a drop in biomass density (Fig. S7a-c) in cells immediately adjacent to the wound, as they underwent epithelial-mesenchymal transition. Previous studies have shown that most of the tissue tension that results in wound closure comes from the motility forces of cells that are distant from the wound^58,59^. The picture that emerges from our results is that cells at the immediate wound edge become motile first and move into the wound, mechanically stretching cells in the rows behind them. As this wave of mechanical stretch propagates deeper into the tissue, it causes a drop in biomass density, resulting in depolarization of membrane potential. This depolarization provides signals to upregulate motility, biomass production and proliferation to enhance wound healing.

### Depolarization enhances efficiency of wound healing

We tested whether modulation of membrane potential directly affects efficiency of wound healing by applying depolarizing and hyperpolarizing drugs and quantifying wound healing speed. In this experiment we used CHO cells instead of MDCK cells because MDCK tissues tended to detach entirely during scratch wounding in our protocol. We found that even low concentrations of hyperpolarizing drugs (valinomycin) dramatically slowed down wound healing (Fig. 3f, lower panel & Fig. 3g, yellow circles) compared to controls. We previously established that increasing external osmolarity results in hyperpolarization of membrane potential (Fig. S4), and this also phenocopies the effect of hyperpolarizing drugs (Fig. 3g, green circles & Fig. S8). Remarkably, we found that the efficiency of wound healing could be increased, becoming even faster than wild-type tissues, by adding depolarizing drugs (Fig. 3f, middle & Fig. 3g, red circles; compare with Fig.3f, upper & Fig. 3g, cyan circles; see also Fig. S for additional drugs). These data indicate that depolarization accelerates the upregulation of motility and proliferation compared to the control tissue, where depolarization is triggered later by stretching.

### Membrane potential depolarization after wounding in *Xenopus* tail regeneration

To test whether membrane potential depolarization due to a decrease in biomass density occurs *in vivo*, we used the Xenopus embryo (stage 40) tail regeneration as a model system (Fig. S9a-b). We observed a gradient of depolarization from the wound edge and spanning over 6 cell layers 10 minutes after tail amputation (Fig. 3i). Using NoRI, we also observed a reduction in biomass density of cells at the wound edge immediately after tail amputation (Fig. 3j top panels). After 10 minutes, we observed that 5 to 6 layers of cells next to the wound edge had become significantly dilute (Fig. 3j bottom panels). This drop in biomass density is probably due to stretching from a rapid wound healing response, where cell layers from the two lateral sides rapidly roll over and merge to seal the cut wound edge. An experiment in the fresh-water polyp *Hydra vulgaris* after amputation of the body trunk showed a very similar result (Fig. S9c). The coordination of growth and wound healing by membrane potential due to changes in biomass density thus appears to be an ancient mechanism that goes back to the earliest metazoan ancestors.

### Membrane potential controls nuclear localization of the Hippo transcription factor YAP

The Hippo signaling pathway is one of the classic mediators of mechano-transduction^60^. The key downstream effector of this pathway is the transcription factor YAP, which shuttles between nucleus and cytoplasm and promotes cell proliferation when localized to the nucleus^60,61^. However, the mechanisms that render Hippo signaling mechano-sensitive are incompletely understood and no physiological signals that activate FAT1, the upstream membrane-bound mediator of Hippo signaling have yet been discovered. We therefore decided to test whether Hippo signaling is directly affected by changes in membrane potential.

Indeed, we observed continuous modulation of nuclear/cytoplasm ratio of YAP by cell number density in our system (Fig. 4a-c), coinciding with the changes in membrane potential and biomass density that we observed (Fig. 1). To test the hypothesis that YAP localization is directly under the control of membrane potential, we quantified YAP localization at different cell densities, in combination with hyperpolarizing and depolarizing drugs. In sub-confluent and low number density cells (which we will call “sparse”, in contrast to fully confluent, high number density “dense” cells) hyperpolarization caused significant nuclear exclusion and cytoplasmic localization of YAP (Fig. 4d and Fig. S10a). Conversely, in dense cells depolarization of membrane potential using a range of drugs with different mechanisms of action was sufficient to cause YAP nuclear reentry (Fig. 4e, and Figs S10b and S11).

**Figure 4:**
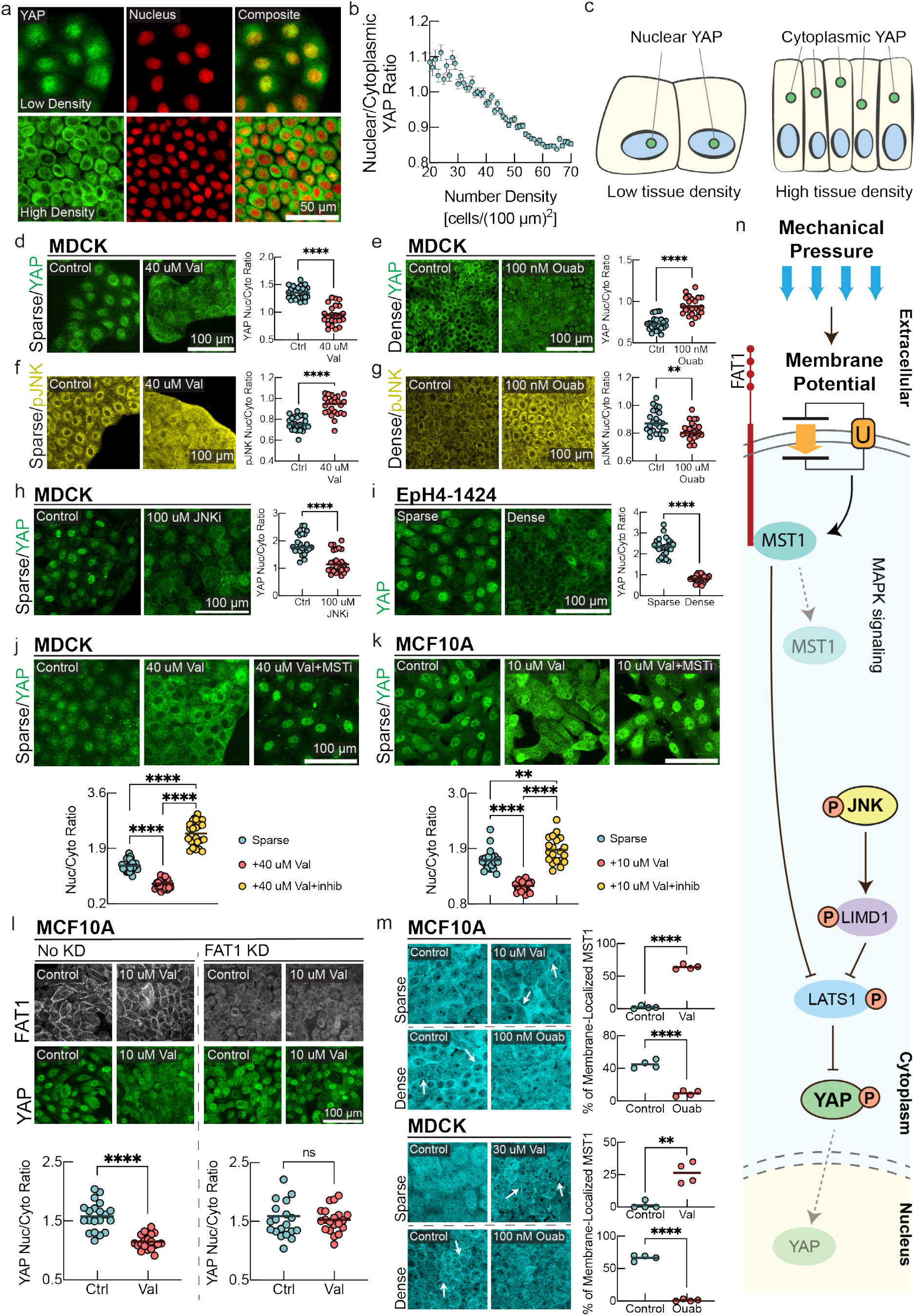
Hippo and MAPK pathways mediate membrane potential signaling. **a**, YAP immunostaining and fluorescent nuclei images in low density (top panel) and high density MDCK monolayers (bottom panel). **b**, Quantification of nuclear/cytoplasmic YAP ratio as a function of cell number density. (mean ± s.d., n = 20337, cells binned together by local number density). **c**, Schematic representation of YAP localization at different cell number densities. **d-g**, Immunostaining of MDCK cells for YAP and phosphorylated JNK (pJNK) under sparse (left panels) and dense (right panels) conditions with corresponding drug perturbations.**d-e**, the nucleus/cytoplasm ratio of YAP is measured. **d**, In Sparse cells, YAP localizes to the nucleus and induction of hyperpolarization significantly promotes nuclear exclusion of YAP. **e**, in dense confluent cells, depolarization by ouabain inhibits nuclear exclusion of YAP as compared to the control. In the plots, the line is at mean, n=24, and p-value is measured with an unpaired t-test and is <0.0001 unless otherwise specified. **f**, pJNK predominantly localized in cytoplasm in sparse MDCK cells. Induction of hyperpolarization by valinomycin promotes increased nuclear localization of pJNK. **g**, In dense confluent cells, depolarization by ouabain promotes nuclear exclusion of pJNK as compared to the control. **h**, JNK inhibition promotes significant nuclear exclusion of YAP in sparse MDCK cells. **i**, In Eph4-1424 cells, YAP nuclear localization follows the canonical density dependence of YAP nuclear-cytoplasmic shuttling. **j-k**, YAP immunostaining of sparse MDCK/MCF10A cells with control, valinomycin, and valinomycin with MST inhibitor (XMU-MP-1) conditions. The p-value between sparse and valinomycin+MSTi conditions is 0.0092. **l**, Sparse MCF10A cells with and without FAT1 knockdown (KD) are stained for FAT1 and YAP under control and valinomycin conditions. The nucleus/cytoplasm ratio for FAT1KD cells under control vs valinomycin has a p-value of 0.558 (ns). **l**, MST1 immunostaining of MCF10A/MDCK cells under sparse control, sparse valinomycin, dense control, and dense ouabain conditions. White arrows highlight areas of increased MST1 organized membrane clusters. For quantifications in panels. **m**, localization of MST1 in individual cells in randomized fields of view were analyzed. The line is at mean, n=4 ROIs, and p-value is measured with an unpaired t-test. The p-value is 0.0012 for MDCK sparse control v. valinomycin and <0.0001 for other plots. **n**, Schematic diagram summarizing signal transduction mechanism mediated by membrane potential. Depolarization of membrane potential activates cytoplasmic pJNK, which activates LIMD1 to inhibit LATS1, causing YAP to translocate to the nucleus. In parallel, hyperpolarization of membrane potential causes MST1 to colocalizes with the c-terminus tail of FAT1, assembling the Hippo signalome. This Hippo ‘on’ state prevents the nuclear translocation of YAP.

The observed regulation of YAP by membrane potential can be rationalized in the context of homeostatic regulation of growth by membrane potential. Low tissue density results in low mechanical pressure and depolarized membrane potential, resulting in localization of YAP to the nucleus, where it promotes cell growth and cell cycle progression. Conversely, high tissue density is reflected in high mechanical pressure and hyperpolarized membrane potential, which results in YAP exiting the nucleus and suppresses proliferation. In Figs. S5&S8, we demonstrated that hyperpolarization leads to a lower steady-state cell number density and negatively affects wound healing efficiency, which is consistent with the anti-proliferative effect of nuclear exclusion of YAP under hyperpolarizing conditions. Moreover, a previous observation^20^ of nuclear reentry of YAP in confluent monolayers upon mechanical stretch^20^ can be mechanistically explained by our model, as stretching causes a drop in biomass density and concomitant depolarization of cellular membrane potential (Fig. 2).

### Membrane potential modulates MAPK signaling, affecting YAP localization

Hippo signaling is known to undergo crosstalk with MAPK signaling via pJNK^62^. Since MAPK pathways have been reported to be mechanoresponsive, we next investigated whether membrane potential directly affected MAPK signaling^63–65^, which could affect YAP localization^62^.

We found that in sub-confluent and low number density cells, pJNK predominantly localizes in the cytoplasm, whereas in high density cells pJNK predominantly localizes in the nucleus (Fig. 4f-g, and Fig. S10c-d). If this is caused by membrane polarization, then inducing hyperpolarization in sparse cells should cause pJNK to translocate to the nucleus, while depolarizing dense cells should cause pJNK to become cytoplasmic. We saw precisely these effects upon addition of a hyperpolarizing drug in sparse cells (valinomycin, Fig. 4f, Fig. S10c) and a depolarizing drug to dense cells (ouabain, Fig. 4g, and Fig. S10d). Similarly, we found that p38 signaling, another downstream effector of MAPK pathway that is involved in mechanical signaling, could also be controlled by changing membrane potential (Fig. S12). It was recently demonstrated that depolarization of the membrane potential can induce K-Ras clustering^63^, and ionic interactions can affect the nanoclustering of EGFR^65^, providing a plausible upstream route to MAPK pathway modulation. Nuclear localization of activated JNK and p38 at high tissue density is expected to contribute to tissue density homeostasis by suppressing proliferation, promoting autophagy, inducing apoptosis, or causing cell extrusion^66–68^, while translocation of these signals out of the nucleus would be expected to promote proliferation.

We next asked whether pJNK was involved in controlling YAP localization by adding a JNK inhibitor (SP600125) to sparse MDCK (Fig. 4h) and EpH4-EV cells (Fig. S10e). This caused significant movement of YAP out of the nucleus. We saw the same effect in dense MCF10A cells (Fig. S10f), indicating that JNK crosstalk contributes to the regulation of YAP localization. However, JNK signaling is not the sole mediator of YAP nuclear localization. We tested this using the mouse breast epithelial cell line EpH4-1424, which expresses a constitutive active form of MEK (MEKDD)^69^, activating JNK via ERK^70^. We confirmed the activation of both pERK and pJNK in this cell line (Fig. S10g). If pJNK signaling entirely controls YAP nuclear localization, then YAP should be retained in the nucleus in EpH4-1424 cells even at high density. Instead, we found that YAP still showed nuclear exclusion in these cells (Fig. 4i).

### Membrane potential controls Hippo signalome assembly via FAT1

Since JNK signaling does not fully explain YAP signaling in response to membrane potential, we investigated the canonical Hippo pathway upstream of YAP. Assembly and activation of the Hippo signalome is dependent on the upstream membrane-bound mediator FAT1^71^. When Hippo is on, MST1 becomes active and phosphorylates LATS1/2, which in turn phosphorylates YAP, inhibiting its nuclear localization. However, no upstream physiological signals that activate FAT1 have yet been discovered. We therefore asked whether changes in membrane potential could directly affect FAT1-mediated assembly of the Hippo signalome complex^71^. First, we added a hyperpolarizing drug (valinomycin) to sparse MDCK and MCF10A cells, in combination with an MST1 inhibitor (XMU-MP-1). Hyperpolarization resulted in nuclear exclusion of YAP, which was reversed by the MST1 inhibitor (Fig. 4j-k). Similarly, FAT1 knockdown resulted in nuclear localization of YAP (Fig. 4l), which could not be overridden by membrane potential hyperpolarization (Fig. 4l). Thus, the nuclear exclusion of YAP that results from hyperpolarization of membrane potential is mediated by MST1, and absence of FAT1 abrogates all effects of membrane potential hyperpolarization on YAP.

To assess if membrane potential directly affected assembly of the FAT1-MST1 signalome complex, we quantified the localization of MST1. In sparse MDCK and MCF10a cells, MST1 is diffuse in the cytoplasm (Fig. 4m). However, in dense cells, when the Hippo signalome is assembled and Hippo is in the ‘on’ state, MST1 is recruited to the membrane (Fig. 4m). We tested whether membrane potential affected this process. It does: hyperpolarization by valinomycin caused MST1 to form organized clusters on the membrane of sparse cells, mimicking the dense state (Fig. 4m, left panel). Conversely, depolarization by ouabain disrupted MST1 membrane localization in dense cells, rendering MST1 diffuse in the cytoplasm (Fig. 4m right panel). These data suggest that membrane potential controls Hippo signalome assembly and could be the upstream physiological signal that enables transduction of mechanical pressure via the Hippo pathway.

## DISCUSSION

Our data indicate that the membrane potential of a cell provides a quasi-instantaneous, globally integrated readout of cytoplasmic density and mechanical pressure that informs the cell of whether growth is needed to maintain tissue homeostasis. The coupling between mechanical pressure, cytoplasmic density and membrane potential is based only on elementary physical laws, requires no active regulation, and is present even in thermodynamic equilibrium^44^. Therefore, even primitive cells, lacking active transport and confined by a simple cell wall, could have utilized membrane potential as a readout of their mechanical state and biomass concentration. Membrane-potential-mediated pressure-transduction may therefore have been among the first homeostatic mechanisms to emerge in multicellular organisms. Note that while our model assumes no active regulation or amplification of membrane potential, it is possible that cells may have evolved to amplify changes of membrane potential signaling in response to mechanical pressure, enhancing their ability to regulate their growth in response to tissue density and mechanical forces. Indeed, the changes we observed in membrane potential in our experiments are sometimes larger than those expected from our model (Fig. S3).

We find that membrane potential controls signaling through the Hippo and MAPK pathways. Our survey of signaling pathways was far from exhaustive, and in principle membrane potential could affect any signaling pathway originating at the plasma membrane, including those involving cadherins, receptors, and voltage-gated ion channels. For example, calcium signaling would be an ideal effector of membrane-potential-mediated transduction of mechanical pressure. This could occur either via voltage-gated channels in the plasma membrane or the strongly voltage-dependent equilibrium calcium concentration that results from the calcium ion’s two positive charges. Indeed, flux through key calcium channels like Piezo has been shown to strongly depend on membrane potential^72^ and Piezo has been shown to be voltage-gated^73^.

Pressure transduction via membrane potential is unlikely to be limited to epithelia. For instance, stem cell differentiation has been shown to depend on both cell volume and osmotic pressure^74,75^, as well as membrane potential^76^, suggesting a role of membrane-potential-mediated mechanical signaling. Membrane potential could also have a role in neuronal mechano-transduction in situations where mechanical forces affect cell volume. This could result in changes in resting membrane potential and trigger voltage-gated channels and calcium signaling^72,73^, independent of tension in the plasma membrane.

Our work shows, for the first time, that cellular biomass density is not constant, but that dramatic changes in biomass density occur in physiological conditions such as wound healing and epithelial maturation. Such changes in biomass density could have other effects beyond their effect on membrane potential. There is growing interest in the possibility that macromolecular condensates may be important in signal transduction. Physical phase transitions that result in condensates are strongly dependent on macromolecular density. Thus, changes in biomass density could directly trigger condensate formation or dissolution, providing additional amplification of signal transduction.

Finally, the importance of membrane potential in tissue growth control suggests that misregulation of membrane potential or its downstream signaling could have dramatic consequences for tissue homeostasis, potentially resulting in tumorigenesis when cells stop responding to the mechanical cues from their environment. This may explain the frequent occurrence of ectopic expression of neuronal ion channels in tumors^77^ and the commonly observed phenotype of depolarized resting membrane potential in cancer^78^.

## Acknowledgements

We would like to thank Becky Ward, Sean Megason, Tim Mitchison, Jan Skotheim, and Xavier Trepat for their comments and feedback on the manuscript. We would like to thank Peter Sorger and Clarence Yapp for access to their high content imaging systems. Thanks to Eugenia Piddini (University of Bristol), J.J. Fredberg and C.Y. Park (Harvard T.H. Chan School of Public Health) for helping us with MDCK cells. We thank Nikon imaging center at Harvard Medical School for help with all fluorescence imaging experiments. Specifically, we thank Jennifer Waters, Tally Lambert, Anna Payne-Tobin Jost for their valuable support with the microscopy. Thanks to Florian Engert for helping us with Hydra experiments, and special thanks to Andrew Murphy of Sean Megason lab for helping us with Hydra husbandry. Thanks to Bill Jia of Megason lab for valuable comments on membrane potential measurement. Thanks to Matthew Yonas (Harvard medical school) for helping with scratch wound assay. This project was supported by a MIRA grant (5R35GM137895) and an HMS Junior Faculty Armenise grant to M.B. N.C.B was supported by following grants National Science Foundation Graduate Research Fellowship Program (DGE 2140743) and Systems, Synthetic, and Quantitative Biology Training grant award (T32GM135014). YH was supported by HCRP of Harvard College. Any opinions, findings, and conclusions or recommendations expressed in this material are those of the author(s) and do not necessarily reflect the views of the National Science Foundation. Leonid Peshkin was supported by R01AG073341 and R24OD031956 and Marc W. Kirschner by R01AG073341 and MIRA R35 GM145248. Christophe Dupre is supported by the Swiss National Science Foundation Early Postdoc Mobility Fellowship (P2SKP3-187684).

## References

1. Zhu, J. & Thompson, C. B. Metabolic regulation of cell growth and proliferation. Nat Rev Mol Cell Biol 20, 436–450 (2019).

2. Hosios, A. M. et al. Amino Acids Rather than Glucose Account for the Majority of Cell Mass in Proliferating Mammalian Cells. Dev Cell 36, 540–549 (2016).

3. Palm, W. & Thompson, C. B. Nutrient acquisition strategies of mammalian cells. Nature 546, 234–242 (2017).

4. Vander Heiden, M. G., Cantley, L. C. & Thompson, C. B. Understanding the Warburg Effect: The Metabolic Requirements of Cell Proliferation. Science (1979) 324, 1029–1033 (2009).

5. Pavlova, N. N., Zhu, J. & Thompson, C. B. The hallmarks of cancer metabolism: Still emerging. Cell Metab 34, 355–377 (2022).

6. Barbazan, J. et al. Cancer-associated fibroblasts actively compress cancer cells and modulate mechanotransduction. Nat Commun 14, 6966 (2023).

7. Kim, S., Uroz, M., Bays, J. L. & Chen, C. S. Harnessing Mechanobiology for Tissue Engineering. Dev Cell 56, 180–191 (2021).

8. Humphrey, J. D., Dufresne, E. R. & Schwartz, M. A. Mechanotransduction and extracellular matrix homeostasis. Nat Rev Mol Cell Biol 15, 802–812 (2014).

9. Pensalfini, M. & Tepole, A. B. Mechano-biological and bio-mechanical pathways in cutaneous wound healing. PLoS Comput Biol 19, e1010902 (2023).

10. Rosińczuk, J., Taradaj, J., Dymarek, R. & Sopel, M. Mechanoregulation of Wound Healing and Skin Homeostasis. Biomed Res Int 2016, 1–13 (2016).

11. Cambria, E. et al. Linking cell mechanical memory and cancer metastasis. Nat Rev Cancer 24, 216–228 (2024).

12. Chaudhuri, P. K., Low, B. C. & Lim, C. T. Mechanobiology of Tumor Growth. Chem Rev 118, 6499–6515 (2018).

13. Papavassiliou, K. A., Basdra, E. K. & Papavassiliou, A. G. The emerging promise of tumour mechanobiology in cancer treatment. Eur J Cancer 190, 112938 (2023).

14. Clevenger, A. J. et al. Advances in cancer mechanobiology: Metastasis, mechanics, and materials. APL Bioeng 8, (2024).

15. Wong, S. H. D. et al. Mechanical manipulation of cancer cell tumorigenicity via heat shock protein signaling. Sci Adv 9, (2023).

16. Li, C. et al. Extracellular matrix-derived mechanical force governs breast cancer cell stemness and quiescence transition through integrin-DDR signaling. Signal Transduct Target Ther 8, 247 (2023).

17. Labernadie, A. et al. A mechanically active heterotypic E-cadherin/N-cadherin adhesion enables fibroblasts to drive cancer cell invasion. Nat Cell Biol 19, 224–237 (2017).

18. Aragona, M. et al. A Mechanical Checkpoint Controls Multicellular Growth through YAP/TAZ Regulation by Actin-Processing Factors. Cell 154, 1047–1059 (2013).

19. Piccolo, S., Dupont, S. & Cordenonsi, M. The Biology of YAP/TAZ: Hippo Signaling and Beyond. Physiol Rev 94, 1287–1312 (2014).

20. Benham-Pyle, B. W., Pruitt, B. L. & Nelson, W. J. Mechanical strain induces E-cadherin– dependent Yap1 and β-catenin activation to drive cell cycle entry. Science (1979) 348, 1024–1027 (2015).

21. Luo, J. et al. The oncogenic roles and clinical implications of YAP/TAZ in breast cancer. Br J Cancer 128, 1611–1624 (2023).

22. Li, H. et al. YAP/TAZ drives cell proliferation and tumour growth via a polyamine–eIF5A hypusination–LSD1 axis. Nat Cell Biol 24, 373–383 (2022).

23. Saotome, K. et al. Structure of the mechanically activated ion channel Piezo1. Nature 554, 481–486 (2018).

24. Ranade, S. S. et al. Piezo2 is the major transducer of mechanical forces for touch sensation in mice. Nature 516, 121–125 (2014).

25. Coste, B. et al. Piezo proteins are pore-forming subunits of mechanically activated channels. Nature 483, 176–181 (2012).

26. Kosmalska, A. J. et al. Physical principles of membrane remodelling during cell mechanoadaptation. Nat Commun 6, 7292 (2015).

27. Shi, Z., Graber, Z. T., Baumgart, T., Stone, H. A. & Cohen, A. E. Cell Membranes Resist Flow. Cell 175, 1769-1779.e13 (2018).

28. Devany, J., Falk, M. J., Holt, L. J., Murugan, A. & Gardel, M. L. Epithelial tissue confinement inhibits cell growth and leads to volume-reducing divisions. Dev Cell 58, 1462-1476.e8 (2023).

29. Oh, S. et al. Protein and lipid mass concentration measurement in tissues by stimulated Raman scattering microscopy. Proc Natl Acad Sci U S A 119, e2117938119 (2022).

30. Buckley, C. E. & St Johnston, D. Apical–basal polarity and the control of epithelial form and function. Nat Rev Mol Cell Biol 23, 559–577 (2022).

31. Park, H. et al. Three-dimensional refractive index tomograms and deformability of individual human red blood cells from cord blood of newborn infants and maternal blood. 10.1117/1.JBO.20.11.111208 20, 111208 (2015).

32. Shin, S., Kim, K., Yoon, J. & Park, Y. Active illumination using a digital micromirror device for quantitative phase imaging. Optics Letters, Vol. 40, Issue 22, pp. 5407-5410 40, 5407–5410 (2015).

33. Shaw, K. L., Grimsley, G. R., Yakovlev, G. I., Makarov, A. A. & Pace, C. N. The effect of net charge on the solubility, activity, and stability of ribonuclease Sa. Protein Science 10, 1206–1215 (2001).

34. Tanford, C. The Interpretation Of Hydrogen Ion Titration Curves Of Proteins. in 69–165 (1963). doi:10.1016/S0065-3233(08)60052-2.

35. Vallina Estrada, E., Zhang, N., Wennerström, H., Danielsson, J. & Oliveberg, M. Diffusive intracellular interactions: On the role of protein net charge and functional adaptation. Curr Opin Struct Biol 81, 102625 (2023).

36. Donnan, F. G. Theorie der Membrangleichgewichte und Membranpotentiale bei Vorhandensein von nicht dialysierenden Elektrolyten. Ein Beitrag zur physikalisch-chemischen Physiologie. Zeitschrift für Elektrochemie und angewandte physikalische Chemie 17, 572–581 (1911).

37. Abdelfattah, A. S. et al. Bright and photostable chemigenetic indicators for extended in vivo voltage imaging. Science (1979) 365, 699–704 (2019).

38. Cone, C. D. & Tongier, M. Contact inhibition of division: involvement of the electrical transmembrane potential. J Cell Physiol 82, 373–86 (1973).

39. Barboiu, M. et al. Polarized Water Wires under Confinement in Chiral Channels. J Phys Chem B 119, 8707–8717 (2015).

40. Barboiu, M. et al. An artificial primitive mimic of the Gramicidin-A channel. Nat Commun 5, 4142 (2014).

41. Goldman, D. E. POTENTIAL, IMPEDANCE, AND RECTIFICATION IN MEMBRANES. J Gen Physiol 27, 37 (1943).

42. Hille, B. Ion channels of excitable membranes. in (2001).

43. Tosteson, D. C. & Hoffman, J. F. Regulation of Cell Volume by Active Cation Transport in High and Low Potassium Sheep Red Cells. Journal of General Physiology 44, 169–194 (1960).

44. Kay, A. R. & Blaustein, M. P. Evolution of our understanding of cell volume regulation by the pump-leak mechanism. Journal of General Physiology 151, 407–416 (2019).

45. Armstrong, C. M. The Na/K pump, Cl ion, and osmotic stabilization of cells. Proceedings of the National Academy of Sciences 100, 6257–6262 (2003).

46. Abdul Kadir, L., Stacey, M. & Barrett-Jolley, R. Emerging Roles of the Membrane Potential: Action Beyond the Action Potential. Front Physiol 9, (2018).

47. Manning, G. S. Counterion Binding in Polyelectrolyte Theory. Acc Chem Res 12, 443–449 (1979).

48. Manning, G. S. Limiting Laws and Counterion Condensation in Polyelectrolyte Solutions I. Colligative Properties. J Chem Phys 51, 924–933 (1969).

49. Manning, G. S. Limiting laws and counterion condensation in polyelectrolyte solutions. Biophys Chem 7, 95–102 (1977).

50. Milo, R., Jorgensen, P., Moran, U., Weber, G. & Springer, M. BioNumbers—the database of key numbers in molecular and cell biology. Nucleic Acids Res 38, D750 (2010).

51. Eisenhoffer, G. T. & Rosenblatt, J. Bringing balance by force: live cell extrusion controls epithelial cell numbers. Trends Cell Biol 23, 185–192 (2013).

52. Eisenhoffer, G. T. et al. Crowding induces live cell extrusion to maintain homeostatic cell numbers in epithelia. Nature 484, 546–549 (2012).

53. Park, J. et al. Screening fluorescent voltage indicators with spontaneously spiking HEK cells. PLoS One 8, (2013).

54. Burgstahler, R. et al. Confocal ratiometric voltage imaging of cultured human keratinocytes reveals layer-specific responses to ATP. Am J Physiol Cell Physiol 284, (2003).

55. Suarez-Arnedo, A. et al. An image J plugin for the high throughput image analysis of in vitro scratch wound healing assays. PLoS One 15, e0232565 (2020).

56. Chifflet, S. et al. Early and late calcium waves during wound healing in corneal endothelial cells. Wound Repair and Regeneration 20, 28–37 (2012).

57. Reid, B. & Zhao, M. The Electrical Response to Injury: Molecular Mechanisms and Wound Healing. Adv Wound Care (New Rochelle) 3, 184–201 (2014).

58. Trepat, X. et al. Physical forces during collective cell migration. Nat Phys 5, 426–430 (2009).

59. Ladoux, B. Cells guided on their journey. Nat Phys 5, 377–378 (2009).

60. Panciera, T., Azzolin, L., Cordenonsi, M. & Piccolo, S. Mechanobiology of YAP and TAZ in physiology and disease. Nat Rev Mol Cell Biol 18, 758–770 (2017).

61. Dupont, S. et al. Role of YAP/TAZ in mechanotransduction. Nature 474, 179–183 (2011).

62. Codelia, V. A., Sun, G. & Irvine, K. D. Regulation of YAP by Mechanical Strain through Jnk and Hippo Signaling. Current Biology 24, 2012–2017 (2014).

63. Zhou, Y. et al. Membrane potential modulates plasma membrane phospholipid dynamics and K-Ras signaling. Science (1979) 349, 873–876 (2015).

64. Zhou, Y., Prakash, P., Gorfe, A. A. & Hancock, J. F. Ras and the Plasma Membrane: A Complicated Relationship. Cold Spring Harb Perspect Med 8, a031831 (2018).

65. Wang, Y. et al. Regulation of EGFR nanocluster formation by ionic protein-lipid interaction. Cell Res 24, 959–976 (2014).

66. Zeke, A., Misheva, M., Reményi, A. & Bogoyevitch, M. A. JNK Signaling: Regulation and Functions Based on Complex Protein-Protein Partnerships. Microbiol Mol Biol Rev 80, 793 (2016).

67. Maik-Rachline, G., Lifshits, L. & Seger, R. Nuclear P38: Roles in Physiological and Pathological Processes and Regulation of Nuclear Translocation. International Journal of Molecular Sciences 2020, Vol. 21, Page 6102 21, 6102 (2020).

68. Dhanasekaran, D. N. & Reddy, E. P. JNK signaling in apoptosis. Oncogene 27, 6245–6251 (2008).

69. Pinkas, J. & Leder, P. MEK1 signaling mediates transformation and metastasis of EpH4 mammary epithelial cells independent of an epithelial to mesenchymal transition. Cancer Res 62, 4781–90 (2002).

70. Lopez-Bergami, P. et al. Rewired ERK-JNK Signaling Pathways in Melanoma. Cancer Cell 11, 447–460 (2007).

71. Martin, D. et al. Assembly and activation of the Hippo signalome by FAT1 tumor suppressor. Nat Commun 9, 2372 (2018).

72. Coste, B. et al. Piezo1 and Piezo2 are essential components of distinct mechanically activated cation channels. Science (1979) 330, 55–60 (2010).

73. Moroni, M., Servin-Vences, M. R., Fleischer, R., Sánchez-Carranza, O. & Lewin, G. R. Voltage gating of mechanosensitive PIEZO channels. Nature Communications 2018 9:1 9, 1–15 (2018).

74. Guo, M. et al. Cell volume change through water efflux impacts cell stiffness and stem cell fate. Proc Natl Acad Sci U S A 114, E8618–E8627 (2017).

75. Fan, Y. L., Zhao, H. C. & Feng, X. Q. Hypertonic pressure affects the pluripotency and self-renewal of mouse embryonic stem cells. Stem Cell Res 56, 102537 (2021).

76. Levin, M. Leading Edge Bioelectric signaling: Reprogrammable circuits underlying embryogenesis, regeneration, and cancer. Cell 184, 1971–1989 (2021).

77. Li, M. & Xiong, Z. G. Ion channels as targets for cancer therapy. Int J Physiol Pathophysiol Pharmacol 3, 156 (2011).

78. Yang, M. & Brackenbury, W. J. Membrane potential and cancer progression. Front Physiol 4, (2013).

