## supplementary_Information for "Membrane potential mediates the cellular response to mechanical pressure"

### Membrane potential as master regulator of cellular mechano-transduction

Mukherjee et al.

### Supplementary Information

#### Table of Contents

**TABLE OF CONTENTS** 1

**SUPPLEMENTARY NOTE 1: MECHANO-ELECTRO-OSMOTIC MODEL OF THE CELL** 2

**ION FLUX BALANCE** 2
**FORCE BALANCE** 3
**CHARGE BALANCE** 4
**HOMEOSTASIS OF BIOMASS DENSITY** 5
**MEMBRANE POTENTIAL AS A READOUT OF MECHANICAL PRESSURE AND BIOMASS DENSITY** 6
**ESTIMATION OF PARAMETERS** 6
CELL VOLUME 6
ESTIMATES OF BIOMASS CHARGE 7

**SUPPLEMENTARY NOTE 2: TISSUE SIMULATION** 7

**MODEL OVERVIEW AND MOTIVATION** 7
**SIMULATION DETAILS** 8
APICO-BASAL DIRECTION IN MONO-LAYERED EPITHELIUM 8
MECHANICAL FORCES 8
BIOLOGICAL PROCESSES 9
PARAMETER CALIBRATION 9
BOUNDARY CONDITIONS 10
TEMPORAL PROGRESSION OF SIMULATION 10

**SUPPLEMENTARY FIGURES:** 11

**MATERIALS AND METHODS:** 23

**SUPPLEMENTARY REFERENCES** 27

### Supplementary Note 1: Mechano-electro-osmotic model of the cell

We formulated a simple model, including the three major cytoplasmic ions, sodium, potassium, and chloride. Nevertheless, the model can easily be extended to incorporate additional ion species without affecting any of the main conclusions of this work. The model is derived from ion flux balance of each ion species across the plasma membrane, charge balance of the cell, and mechanical force balance. The major differences as compared to the derivation of the Goldman Hodgkin Katz equation[1] is that the system of equations is closed using mechanical force balance, without assuming a vanishing sum of fluxes of different ions, which is a convenient, yet unphysical assumption. Instead, we assume that passive flux of each ion (due to diffusion and electrophoretic mobility) is balanced by a constant ion-specific rate of active transport. We assume active transport of ions, but no active regulation. When this model is incorporated into a tissue simulation, we assume that membrane potential regulates cellular decision like growth, apoptosis, and motility, but membrane potential, ion transport etc. are not regulated in the model.

#### Ion flux balance

Concentration gradients lead to diffusion flux according to Fick's law and membrane potential $U$ gives rise to an electric field that results in a flux from electrophoretic mobility, according to the Stokes-Einstein relation. Following the derivation of the Goldman equation[1], assuming a constant electric field across a pore of thickness $L$ given by $U/L$, this results in a differential equation relating the flux $j_I$ of ion $I$ (number of ions crossing the membrane per time and per area) and the concentration $[I]$ as a function of position across the membrane denoted by $z$:

$$j_I = -D_I \left( \frac{d[I]}{dz} - \frac{nF}{R_G T} \frac{U}{L} [I] \right), \qquad [1]$$

where $D_I$ is the diffusion constant, $T$ the temperature and $R_G$ and $F$ are the gas and Faraday constants, respectively. The ion charge is denoted by $n$ ( +1 for $Na^+$, $K^+$; -1 for $Cl^-$). Integration over the thickness $L$, with boundary conditions of the intracellular concentration $[I]_{in}$ and extracellular concentration $[I]_{out}$ results in a solution for the flux, that is given by

$$j_I = p_I n\mu \frac{[I]_{in} e^{n\mu} - [I]_{out}}{e^{n\mu} - 1}, \qquad [2]$$

where $\mu = FU/R_G T$ is proportional to the membrane potential and $p_I = D_I/L$ is the permeability. The classical Goldman equation would be obtained from Eq. 2 with the simplifying assumption that the sum of the electric fluxes from all the different ions vanishes. However, in reality, in steady-state each one of these fluxes integrated over the surface of the cell must vanish individually or be compensated by active transport. Assuming homogeneous membrane potential and uniform intracellular concentrations, we can write

$$J_I^{act} = P_I n\mu \frac{[I]_{in} e^{n\mu} - [I]_{out}}{e^{n\mu} - 1}, \qquad [3]$$

where $J_I^{act}$ is the total active transport and $P_I$ is the total permeability over the surface of the cell. Assuming that plasma membrane phospholipid bilayer is impermeable to ions[1], total permeability and total transport activity is determined by the expression level of channels and transporters. Assuming the cell expresses constant levels of pumps and channels as a fraction of total protein, denoted by $\phi_I^{act}$, $\phi_I^{per}$, then both total transport and total permeability scale like the total mass of the cell $M$. In this case, we have $J_I^{act} \sim \phi_I^{act} M$ and $P_I \sim \phi_I^{per} M$ and cell mass $M$ cancels from the equation and we then arrive at a version of Eq. [3] that is independent of the cell mass

$$\phi_I^{act} = \alpha_I \phi_I^{per} n\mu \frac{[I]_{in} e^{n\mu} - [I]_{out}}{e^{n\mu} - 1}, \qquad [4]$$

with all proportionality constants lumped into an effective function $\alpha_I$. Note that $\alpha_I$ can be a proportionality constant but can also depend on other factors. In particular, for the $Na^+/K^+$-ATPase, we assume that its transport activity obeys Michaelis-Menten kinetics dependent on intracellular sodium concentration, i.e. $\alpha_{Na,K} \sim \frac{(Km_{Na})^3 + ([Na]_{in})^3}{([Na]_{in})^3}$. The sodium concentration is taken to the third power because of the stoichiometry of transport by the $Na^+/K^+$-ATPase. We assume that external potassium concentration is relatively high and therefore never limiting. Note that none of the main conclusions from the model depend on these more detailed assumptions and it is straight-forward to modify the model for different active transport fluxes.

Eq. [4] must be applied to each major cellular ion. With the simplifying assumption that active ion transport is largely due to the activity of the $Na^+/K^+$-ATPase, we then obtain the following system of equations

$$-\phi_{ATP} = \alpha_{Na} \phi_{Na}^{per} \mu \frac{[Na]_{out} - [Na]_{in} e^{\mu}}{1 - e^{\mu}}, \qquad [5a]$$

$$\frac{2}{3} \phi_{ATP} = \alpha_K \phi_K^{per} \mu \frac{[K]_{out} - [K]_{in} e^{\mu}}{1 - e^{\mu}}, \qquad [5b]$$

$$0 = -\alpha_{Cl} \phi_{Cl}^{per} \mu \frac{[Cl]_{in} e^{-\mu} - [Cl]_{out}}{e^{-\mu} - 1}. \qquad [5c]$$

Here, $\phi_{ATP}$ denotes the proteome fraction of the $Na^+/K^+$-ATPase, which exports 3 sodium ions for each 2 potassium ions it imports.

#### Force balance

Even small differences between internal and external ion concentrations give rise to a substantial osmotic pressure $p$:

$$p = \left( ([Na]_{in} - [Na]_{out}) + ([K]_{in} - [K]_{out}) + ([Cl]_{in} - [Cl]_{out}) \right) R_G T. \qquad [6]$$

For the cell to neither swell nor shrink, mechanical force balance dictates that this osmotic pressure must be balanced by the sum of other forces on the cell, including forces from cellular structures like the cytoskeleton and forces from the tissue level like extracellular matrix or other cells in the tissue.

On the other hand, according to Eq. [5] ion concentrations are uniquely determined by membrane potential and Eq. [5] can be used to eliminate ion concentrations from Eq. [6]. Hence, we find a one-to-one relation between membrane potential and sum of mechanical forces on the cell (given by the imposed pressure $p$) (Fig. SN1). According to this relationship, with increasing mechanical pressure the cell becomes increasingly negatively polarized. Thus, we argue that membrane potential provides the cell with an elegant, instantaneous way to measure bulk mechanical pressure or tension.

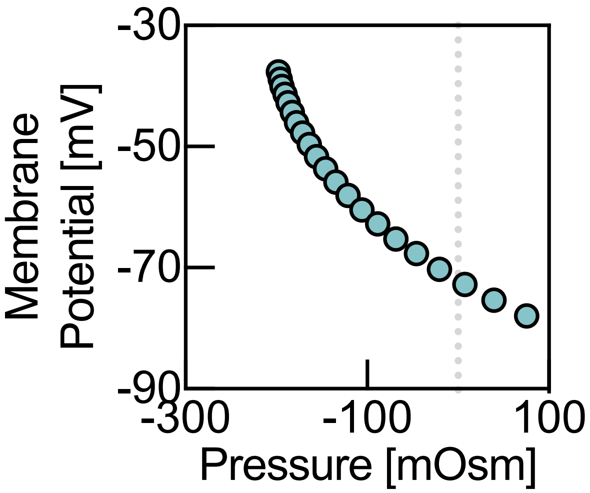


***Fig. SN1: One-to-one mapping between mechanical pressure and membrane potential.*** *According to the model, there is a direct relationship between mechanical pressure applied to the cell and membrane potential.*

We note that it is possible to add additional terms to Eq. [6] to account for the osmotic pressure directly from cellular macromolecules and metabolites. One might assume that part of this osmotic pressure has a constant, density-independent component given by $m_0$, as well as a component proportional to cellular biomass density, given by $m_1\rho$, where $\rho$ is the biomass density of the cell, defined as $\rho = M/V$. This leads to a generalized version of Eq. [6]:

$$p = \big(([Na]_{in} - [Na]_{out}) + ([K]_{in} - [K]_{out}) + ([Cl]_{in} - [Cl]_{out}) + m_1\rho + m_0\big)R_G T. \qquad [6']$$

None of the major conclusions are altered by using Eq. [6'] vs. Eq. [6]. In the simulation, based on our estimates, we assume that the osmotic contribution directly from biomass macromolecules is small and therefore set $m_1$ to zero. The density-independent component given by $m_0$ emerges from metabolite pools and other biomass independent components. This constant osmotic component can be absorbed into a background constant of external pressure in the simulation, bringing us back to Eq. [6].

#### Charge balance

Membrane potential reflects mechanical pressure, but Eqs. [5], [6] are independent of the cell volume (cell volume does enter Eq. [6'] via the biomass density $\rho$, but its contribution is small). This raises the question how cell volume is determined or more specifically how cell volume is coupled to cellular biomass. The answer becomes clear by considering the final physical constraint, closing this system of equations, which is charge balance.

The cell can be thought of as a capacitor and membrane potential arises from net charge distributed over the surface of the cell. Net charge arises on the one hand from summing up charges from intracellular ion concentrations determined by Eq. [5]. The other major component is the net charge of the macromolecular content of the cell, mainly proteins, RNA, and lipids, which carry a net negative charge at physiological pH to aid their solubility[2]. We denote the average net negative charge per cell mass by $c$ and assume a constant biomass composition. Together, we then obtain an equation for membrane potential

$$U = \beta \frac{([Na]_{in}+[K]_{in}-[Cl]_{in})V-cM}{A}, \qquad [7]$$

where $\beta$ is a proportionality constant that resembles an inverse capacitance, $A$ is the surface area of the cell, $V$ the cell volume. This equation dramatically simplifies when considering realistic values for cellular capacitance[1]. Cellular capacitance is so small (i.e. $\beta$ is so large ) that the right side of Eq. [7] is effectively zero. Thus Eq. [7] imposes charge neutrality of the cell, despite non-vanishing membrane potential. Eq. [7] can thus be rewritten as

$$0 = [Na]_{in} + [K]_{in} - [Cl]_{in} - c\rho. \qquad [8]$$

Note that in this equation unlike in Eq. [7], geometry can be completely neglected and only the average biomass density $\rho$ enters the equation.

#### Homeostasis of biomass density

Eqs. [5] & [6] uniquely determine intracellular ion concentrations and membrane potential under an imposed external pressure. Eq. [8] then determines the cell volume for a given cell mass $M$. Cell volume (or biomass density) is determined by the charge concentrations of cellular macromolecules acting on membrane potential. If the cell is compressed, membrane potential becomes more negative, which leads to higher intracellular ion concentrations and an increase in osmotic pressure counteracting this compression. On the other hand, if the cell is stretched to larger sizes, membrane potential becomes less negative and a lower pressure results from lower ion osmolarities. Tension can arise because of active export of ions, e.g. via sodium-potassium-ATPase. This is how the cell achieves homeostasis of biomass density. As the cell produces more biomass, the buildup of macromolecule density leads to an increase of negative charge attracting counterions that result in osmotic pressure. The cell then expands, decreasing the concentration of macromolecules and ions until pressure is balanced. No active regulation is required to achieve this. Fig. SN2 shows pressure as a function of cell volume for two different cell masses.

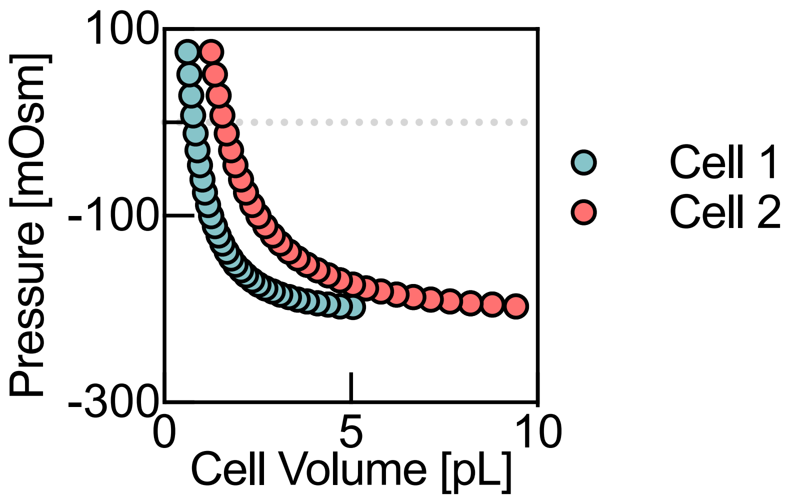


***Fig. SN2: Pressure as a function of cell radius for two different cell masses.*** *Cell 2 has double the biomass of cell 1. More biomass results in mechanical equilibrium (identical pressure) at a higher cell volume.*

To decrease the pressure resulting from intracellular concentrations of counterions and also from other cellular osmolytes, cells actively export ions. The net effect of the $Na^+/K^+$-ATPase is to move ions out of the cell: 3 sodium ions are pumped out for each 2 potassium ions that are pumped into the cell, resulting in a net reduction of intracellular osmolarity. The intracellular concentration of potassium is typically high, and the resulting concentration gradient favors potassium efflux by diffusion, contributing to a reduction of osmotic pressure. Finally, the negatively polarized membrane potential generated by the $Na^+/K^+$-ATPase drives negatively charged $Cl^-$ ions out of the cell, which further decreases intracellular osmotic pressure. Hence, we see that the main function of the $Na^+/K^+$-ATPase is to reduce osmotic pressure and to prevent uncontrolled swelling of the cell.

#### Membrane potential as a readout of mechanical pressure and biomass density

We conclude that membrane potential directly reflects cellular biomass density and total mechanical pressure on the cytoplasm. This pressure results from the combination of cellular forces like cytoskeletal contractility and tissue level forces from other cells or from extracellular matrix. Therefore, using membrane potential, the cell obtains an instantaneous, globally integrated readout of its biomass density and mechanical forces acting upon it. These mechanical forces can result from spatial and physical constraints like cell crowding in tissues that have important implications for cellular growth decisions.

#### Estimation of parameters

##### Cell volume

Highly confluent MDCK cells reach about 70 cells per (100μm)x(100μm) area. We estimate the height of the cell monolayer to be 6.5μm based on our own measurements. Combining these numbers, we obtain an estimate for cell volume at high confluence of ~930μm$^3$.

##### Estimates of biomass charge

To estimate the upper range for concentrations of cellular biomass charge, we first need to estimate the relative abundance of cellular biomass components. We use an estimate of ~30pg RNA per cell (BioNumber ID: 111205)[3] and ~6pg DNA per cell (BioNumber ID: 111206)[3]. Each nucleotide of RNA and DNA is negatively charged and using our estimate of cell volume for confluent epithelial monolayers, we obtain a negative charge concentration of ~97 mM for RNA and ~19mM for DNA.

Assuming 400 amino acids as the mean size of a protein[4] and using frequencies of negatively and positively charged amino acids[5], we estimate a net protein charge of -13*e*. Our NoRI measurements for MDCK cells at high confluence give us a protein density of 0.15g/ml (Fig. 1, main text), resulting in a protein concentration of 3.4mM. This corresponds to a negative charge concentration of ~44mM.

Charge concentration from all remaining biomass components including from lipids and metabolites is difficult to assess. We assume an additional negative charge concentration of ~20mM from all remaining cellular components.

Combining these numbers, we obtain a total negative charge concentration of ~180mM at a biomass density of roughly ~0.15g/ml. This corresponds to ~1.2mole/kg negative biomass charge.

#### Supplementary Note 2: Tissue simulation

##### Model Overview and Motivation

To test more rigorously if our mechano-electro-osmotic model can explain and predict tissue level phenotypes, we built a multi-scale computer simulation, bridging the gap between the physical properties of the cytoplasm, mechanical forces, and cellular signal transduction, which we propose to be regulated by membrane potential.

Mechanical pressure, ion concentrations, cell biomass, cell volume, cell shape, cell position, and resting membrane potential are dynamic, interacting variables that are coupled by elementary physical laws, including those outlined in Supplementary Note 1: Mechano-electro-osmotic model of the cell. We assume that key cellular decisions like growth, apoptosis and cell motility are regulated by the readout of membrane potential, which closely reflects the physiological variables tissue density and mechanical pressure.

In our model, we assume no active regulation of the expression levels or activity of pumps and channels. Only constant expression levels as a fraction of biomass and Michaelis-Menten kinetics of transport activity of sodium-potassium-ATPase based on known biochemical stoichiometry. We make this choice for three reasons: First, for simplicity of the model to limit the number of parameters in the simulation. Second, we want to illustrate that no fine-tuning of model parameters is required for the regulatory mechanism that we propose. Third, we want to illustrate that tissue homeostasis regulated by membrane potential works even in ancient organisms that lack more sophisticated regulatory mechanisms. We have the first multicellular organisms in mind or even single-celled organisms requiring mechano-sensing in their ecological niche. In fact, even in systems without active transport in thermodynamics equilibrium, membrane potential would reflect biomass density and mechanical pressure, which is known as the Gibbs-Donnan effect [2].

Despite these simplifying assumptions one can easily imagine-and in fact, we think it is highly likely-that this ancient mechanism was quickly adapted and refined. For example, voltage-dependent channels and active transport could easily serve to amplify changes in membrane potential, facilitating signal transduction and maximizing signal-to-noise ratio. Indeed, our data indicate that changes in membrane potential with tissue density and pressure are even larger than those expected from our naïve, purely passive model of the cell.

#### Simulation Details

##### Apico-basal direction in mono-layered epithelium

Most epithelial tissues are monolayers. Therefore, we chose a 2-dimensional simulation. In our 2D simulation (in x and y), we do not explicitly model the apico-basal direction (z-direction) and the actin cytoskeleton. We assume a constant height $h$ of the epithelium in the apico-basal direction that might be determined by cytoskeletal elements like microtubules. Cytoplasmic osmotic pressure acting on the apical surface of the cells must be balanced by cytoskeletal contractility, and this contractility could be stabilized by more rigid cytoskeletal structures like apico-basally oriented microtubules.

##### Mechanical forces

Cells positions are represented by their center coordinates in x and y. Shape and volume of cells are determined by a Voronoi construction (scipy.spatial). The mechano-electro-osmotic model, outlined in Supplementary Note 1, is solved for every cell at each timestep based on the cell volume determined by the Voronoi construction and the biomass of the cell. Forces on the cells are then calculated via a finite-difference gradient of an energy function:

$$Energy\ per\ cell\ = P * V + tension * (perimeter - perimeter_0)^2. \qquad [9]$$

Inspired by previous tissue simulations[6], we assume cortical tension modeled by a constant tension and a preferred $perimeter_0 = 3.81\sqrt{area}$, which is right at the jamming transition.

In addition, the simulation includes forces with the substrate $\overrightarrow{f_{sub}}$, given by the sum of dissipative friction, opposing cell motion and motility forces $\overrightarrow{f_{mot}}$,

$$\overrightarrow{f_{sub}} = -\gamma\vec{v} + \overrightarrow{f_{mot}}, \quad [10]$$

where $\vec{v}$ is the velocity of the cell and $\gamma$ is a friction coefficient. For simplicity, we set $f_{mot}$ to zero in our simulation.

#### Biological processes

Biological processes can be implemented and coupled to membrane potential in many ways. Qualitatively, different implementations do not change the results of our simulation. Therefore, we tried to choose the simplest implementation with the key ingredients needed to recapitulate our experimental findings.

*Biomass degradation* occurs at a constant rate, with a half-life $T_{1/2}^{BM} = 5$d.

*Biomass production* has a threshold-linear dependence on membrane potential $U$, when the cell is depolarized below a critical potential:

$$J_{BM} = \xi_{BM}(U - U_0)\theta(U - U_0), \quad [11]$$

where $\theta$ is the heaviside function and $\xi_{BM}$ is a proportionality constant and the threshold membrane potential is denoted by $U_0 = -70$mV. Note that for more polarized cells, membrane potential $U$ is more negative.

*Cell division* is implemented as an "adder"-model, where each cell divides mechanistically after adding a constant critical mass $M_{add}$ since its previous division, based on recent findings [7]. To introduce stochasticity and prevent artifacts from synchronization, the value of $M_{add}$ is drawn from a normal distribution with a mean $\mu_M = 200$pg and a standard deviation $\sigma_M = 50$pg. The mass is divided symmetrically between the mother and daughter cell and the daughter cell is placed 0.1μm in a random direction from the mother cell.

*Cell death* occurs by cells being randomly chosen for apoptosis. The apoptosis rate has a very simple dependence on membrane potential with a constant rate, given by a half-life time of 5 days, and twice this rate when the cell is more polarized than the critical potential $U_0 = -70$mV:

$$T_{1/2}^{apop} = \begin{cases} 5d & for \quad U_0 \leq U \\ 2.5d & for \quad U_0 > U \end{cases}. \quad [12]$$

Cells that undergo apoptosis are simply removed from the simulation.

#### Parameter calibration

Parameters were chosen based on the estimates from Supplementary Note 1. Steady state membrane potential (U*), ion pumping rates and membrane permeability are fully determined by assuming steady-state internal ion concentrations, given by $[Na]_{in}$=5mM, $[K]_{in}$=190mM, $[Cl]_{in}$=15mM, while assuming that the ion concentrations in the medium are given by $[Na]_{out}$=155mM, $[K]_{out}$=5mM, $[Cl]_{out}$=160mM, based on our media composition. $Km_{Na}$ is chosen to be 10mM in the simulation. The remaining parameter $\xi_{BM}$ was determined by choosing the critical potential $U_0$= -70mV and solving the biomass production for steady-state (taking biomass-loss by apoptosis also into account). Note that we confirmed that none of our major results critically depend on these specific parameters.

#### Boundary conditions

Most simulations were done with periodic boundary conditions on an area of 200x200µm to mimic a section of a larger bulk tissue.

#### Temporal progression of simulation

At each timestep of the simulation (dt=0.05h), the time update proceeds in the following steps:

1. Calculating a Voronoi tessellation and use it to determine areas, perimeters, and cell neighbors.
2. Iteratively solve the mechano-electro-osmotic model (see Supplementary Note 1) for each cell, i.e. calculate internal ion-concentrations $[Na]_{in}$, $[[K]_{in}$, $[Cl]_{in}$, membrane potential and pressure.
3. Calculate forces as finite-difference gradients of the effective energy functional given by Eq. [13].
4. Positions are updated by a simple Euler-step.
5. Voronoi tessellation is calculated anew.
6. Cell mass is updated according to biomass production rate (Eq. [15]) minus the constant biomass degradation rate.
7. Each cell has a random chance to undergo apoptosis according to their respective apoptosis rate, given by their half-life (Eq. [16]). If the cell undergoes apoptosis, it is simply removed from the simulation.
8. Cells divide, as described above, if they added a critical mass since the last division, which is drawn from a normal distribution after division.

#### Supplementary Figures:

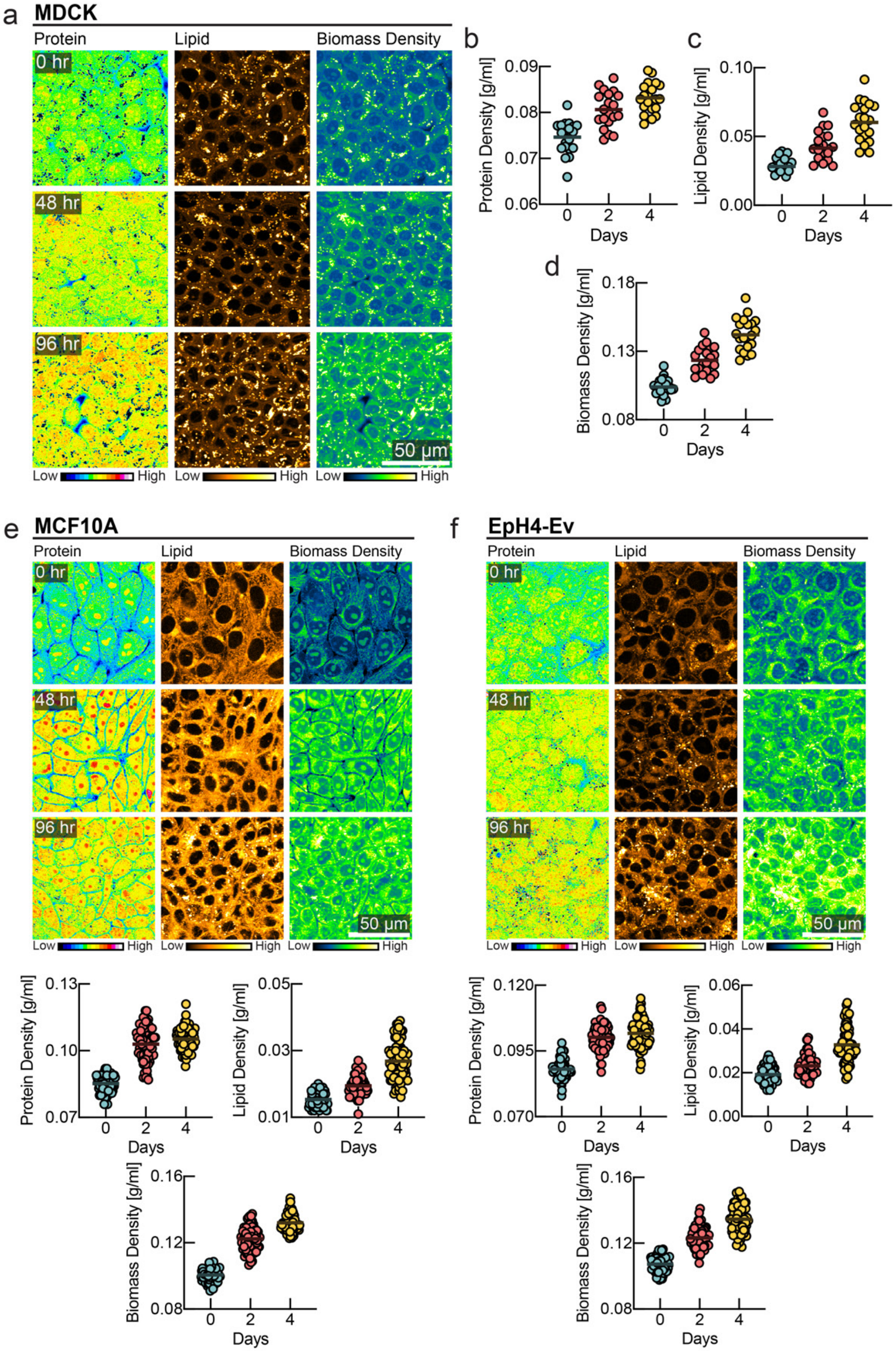


***Fig. S1: Increasing intracellular biomass density with increasing cell number density. a,*** *NoRI images of MDCK epithelial monolayer over 4 days after initial confluence showing contribution of intracellular protein and lipid towards total biomass density.* ***b-d****, Quantification of protein, lipid and biomass density. (line at mean, n=20).* ***e-f,*** *NoRI images of MCF10A/EpH4-Ev monolayers with corresponding quantifications of protein, lipid, and total biomass density. In all tissue types, we observed an increase in protein, lipid, and biomass density across different levels of confluence.*

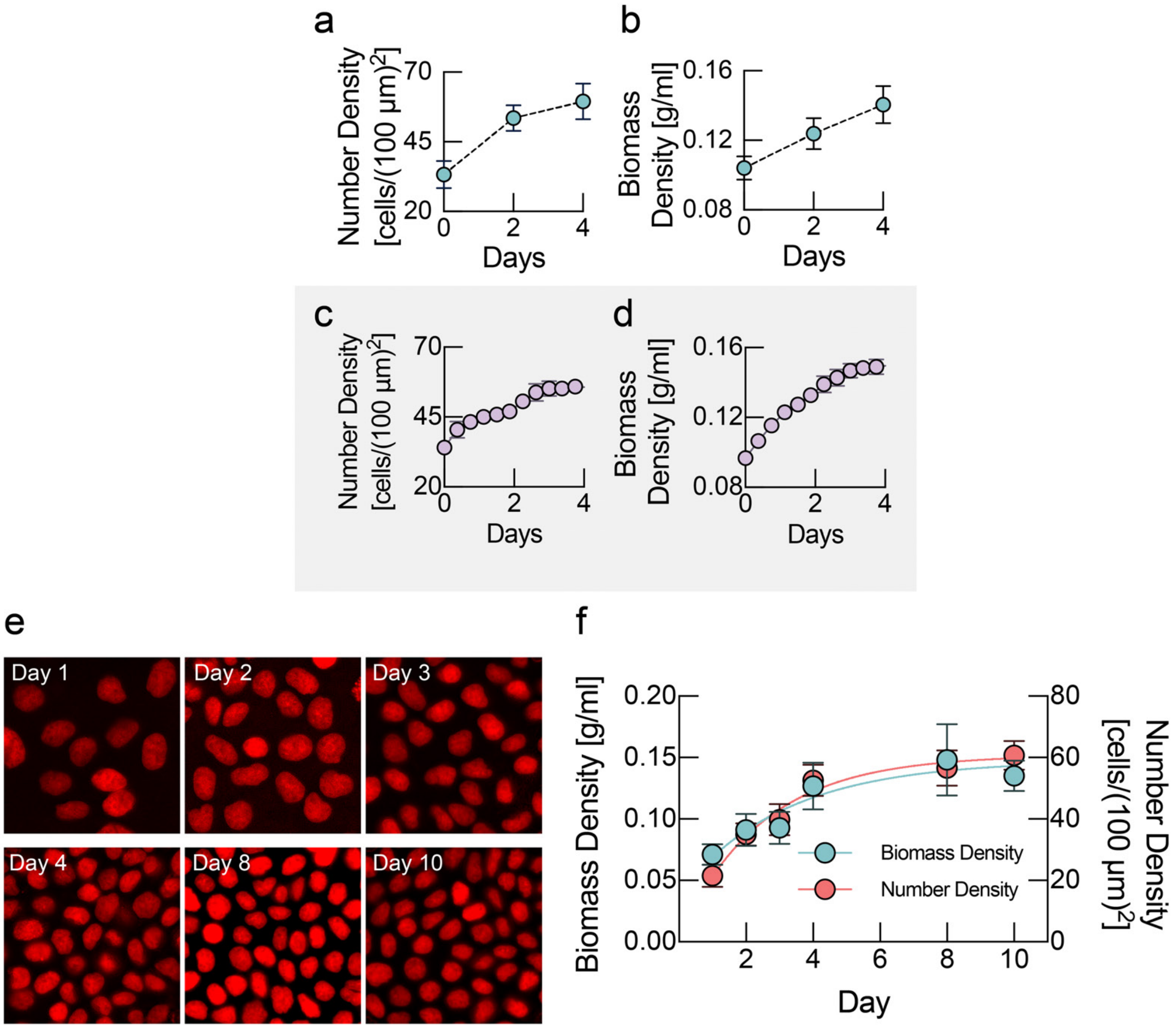


***Fig. S2: Increasing intracellular biomass density and increasing cell number density as a function of time. a,*** *Cell number density from quantified from MDCK monolayers, plotted as function of time. (mean ± s.d., n = 8).* ***b,*** *Cellular biomass density, quantified from MDCK monolayers, increases on the same timescale as cell number density, shown in panel a. (mean ± s.d., n = 15).* ***c-d,*** *Corresponding results from tissue simulation (mean ± s.d., n = 4 simulations).*
***e-f,*** *Cellular biomass density measured by holotomographic microscopy (Tomocube) with increasing cell number density. Left panel shows increasing nuclei density over time in a 75 micron$^2$ field of view. Right plot shows number density (red, mean ± s.d., n=10) and cellular biomass density (blue, mean ± s.d., n=~20) measured using holotomographic microscopy (Tomocube) as a function of time.*

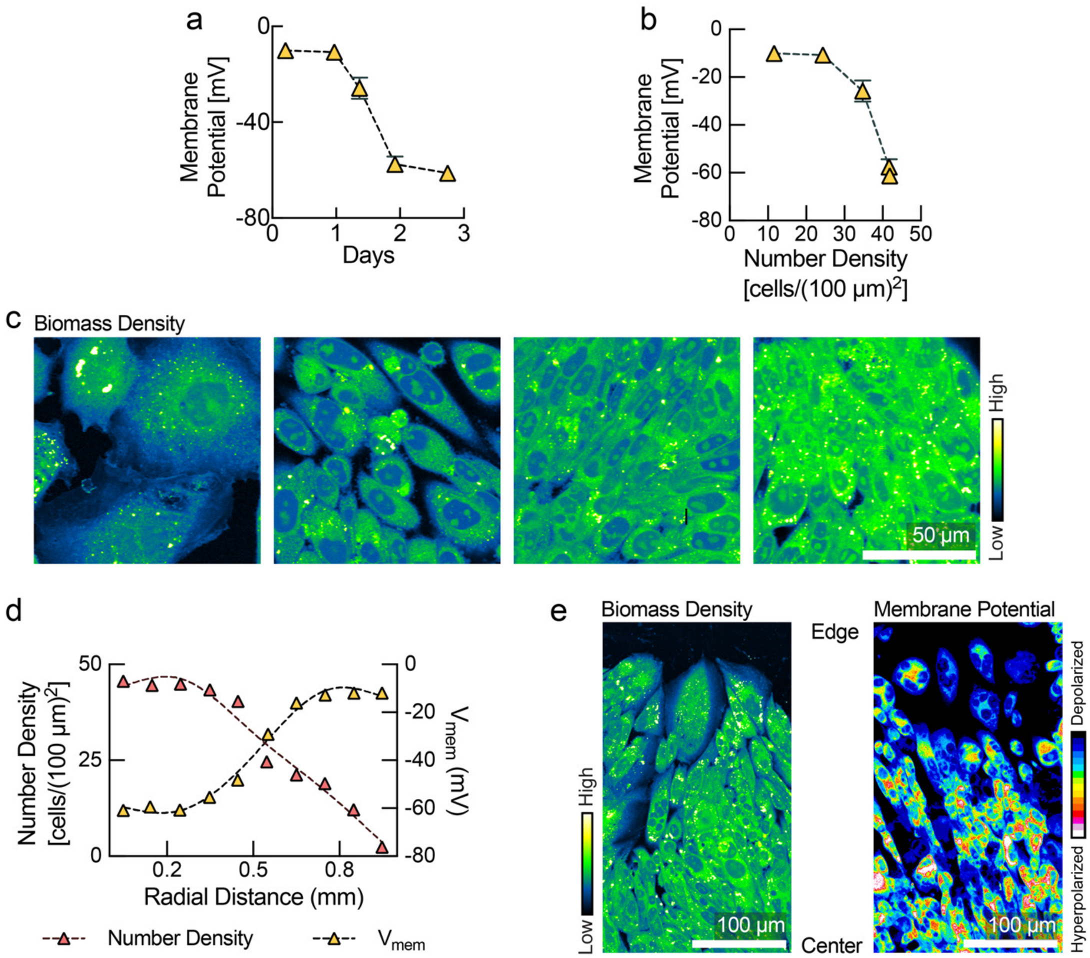


***Fig. S3: CHO K1 cells hyperpolarize with increasing cell number density. a,*** *CHO K1 cells in a growing tissue colony become progressively hyperpolarized with time (replotted from Cone et. al.[8])* ***b,*** *Membrane potential as a function of cell number density in a growing colony of CHO K1 cells (replotted from Cone et. al.[8])* ***c,*** *NoRI images of CHO K1 cells at varying levels of confluence.* ***d,*** *Cell number density (red triangles) and membrane potential (yellow triangles) measured in an expanding colony of CHO K1 cells, as a function of distance from the colony center (replotted from Cone et. al.[8]). Radial distance is measured from the center of the colony. Cells at the colony center are more confluent and hyperpolarized as compared to cells at colony edge.* ***e,*** *Left panel shows NoRI image of a CHO K1, and right panel shows membrane potential measurement by DiOC$_2$(3). Cells at colony edge are more dilute and depolarized as compared to cells in colony center.*

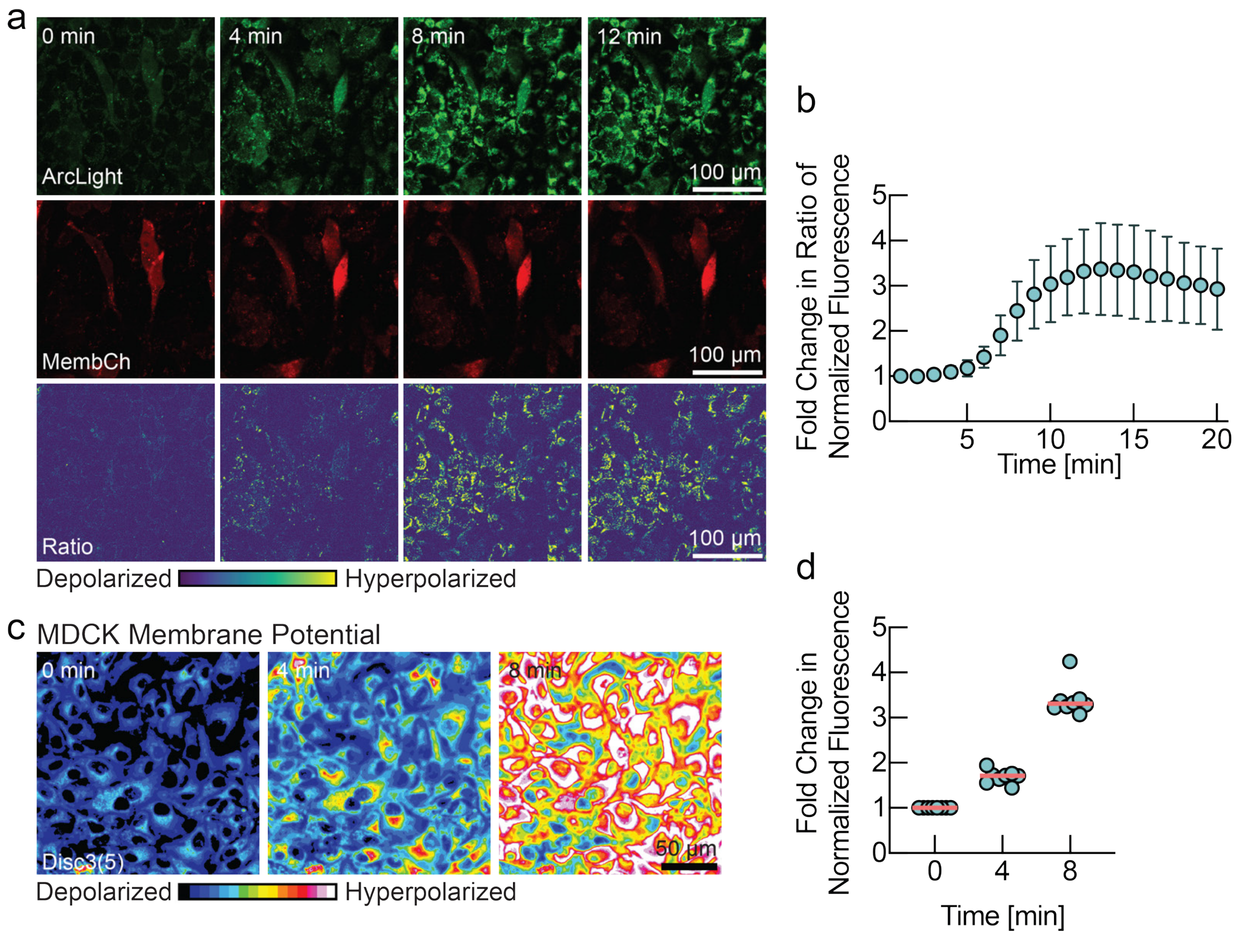


***Fig. S4: CHO K1 and MDCK cells hyperpolarize in response to osmotic pressure. a,*** *CHO K1 cells expressing membrane potential biosensor Arclight and membrane localized mCherry expressed from a bi-cistronic construct. Cells were challenged with 500 mM sucrose to induce osmotic compression. Increase in fluorescence intensity of Arclight channel signifies hyperpolarization of membrane potential. After osmotic compression, cells exhibited an increase in the ratio of Arclight to membrane cherry within minutes of application of osmotic pressure, which continued for 10 minutes before plateauing.* ***b,*** *Fold change of fluorescence ratio quantified from panel a (mean ± s.d., n=8, each cell normalized by initial value).* ***c,*** *MDCK cells were loaded with the membrane potential sensitive dye $DiSC_3(5)$ and then challenged with osmotic pressure. Increase in fluorescence intensity indicates hyperpolarization of membrane potential.* ***d,*** *Fold change of fluorescence intensity quantified from panel c (line at mean, n=8).*

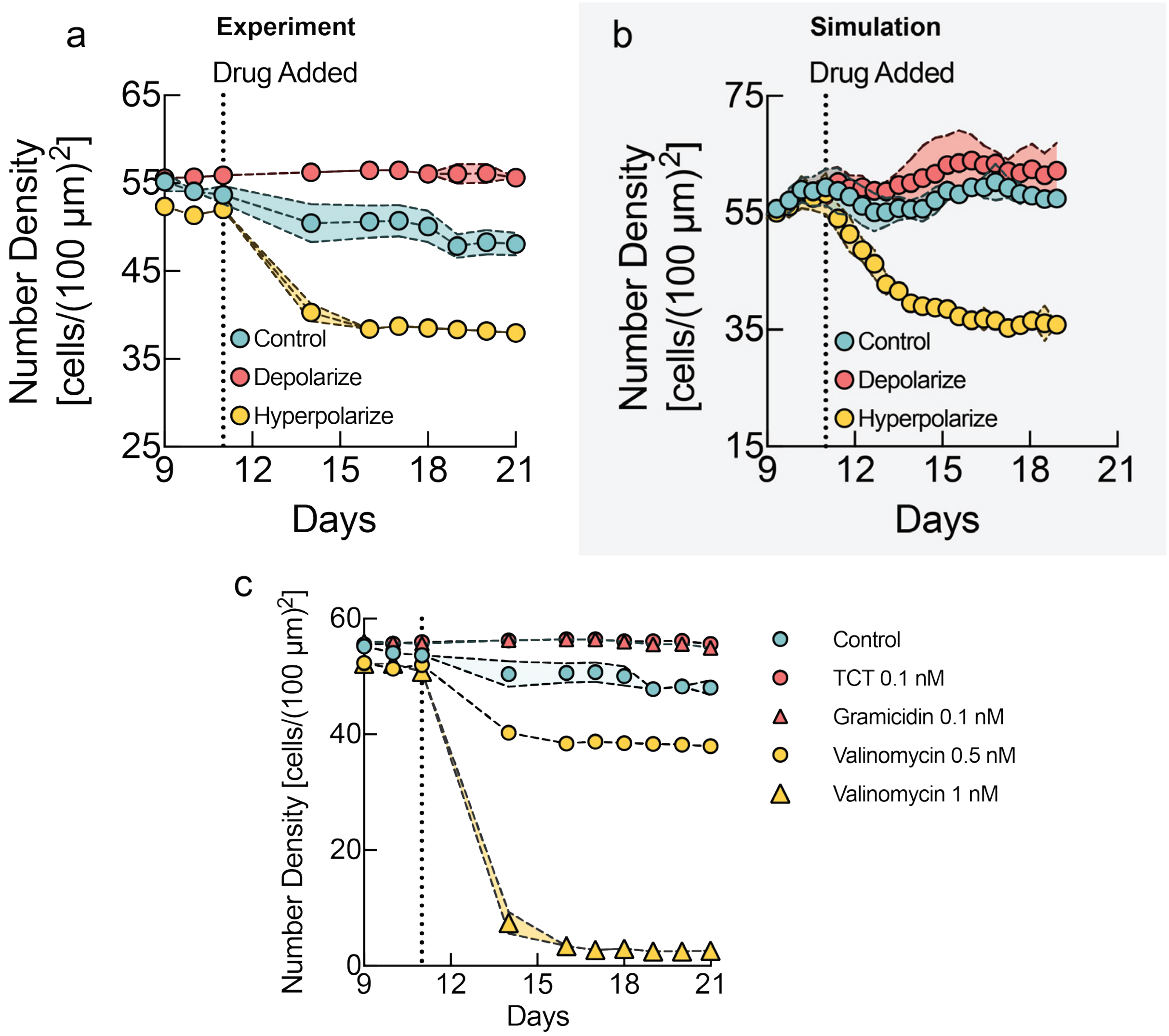


***Fig. S5: Homeostatic cell number density can be modulated with membrane potential drugs**. **a**, Quantification of cell number density (mean ± s.d., n=10 for control, n=3 for drugs) for MDCK cells grown to maximum tissue density and treated with depolarizing (red, TCT) and hyperpolarizing (yellow, valinomycin) drugs as a function of time. Depolarization results in higher tissue density as compared to control. Conversely, hyperpolarization results in a substantially lower tissue density. **b**, Results from the tissue simulation corresponding to experiment in panel a (mean ± s.d., n=4 experiments) **c**, Mild depolarization with a low dose of gramicidin (0.1nM, red triangles) or the gramicidin analog TCT (0.1 nM, red circles) show a similar effect, resulting in a higher cell number density as compared to the control (cyan circles). Cell number density was quantified over time (mean ± s.d., n=10 for control, n=3 for drugs). The hyperpolarizing drug, valinomycin, resulted in a lower steady-state cell number density compared to control, in a dose dependent manner (0.5 nM, yellow circles, and 1nM, yellow triangles).*

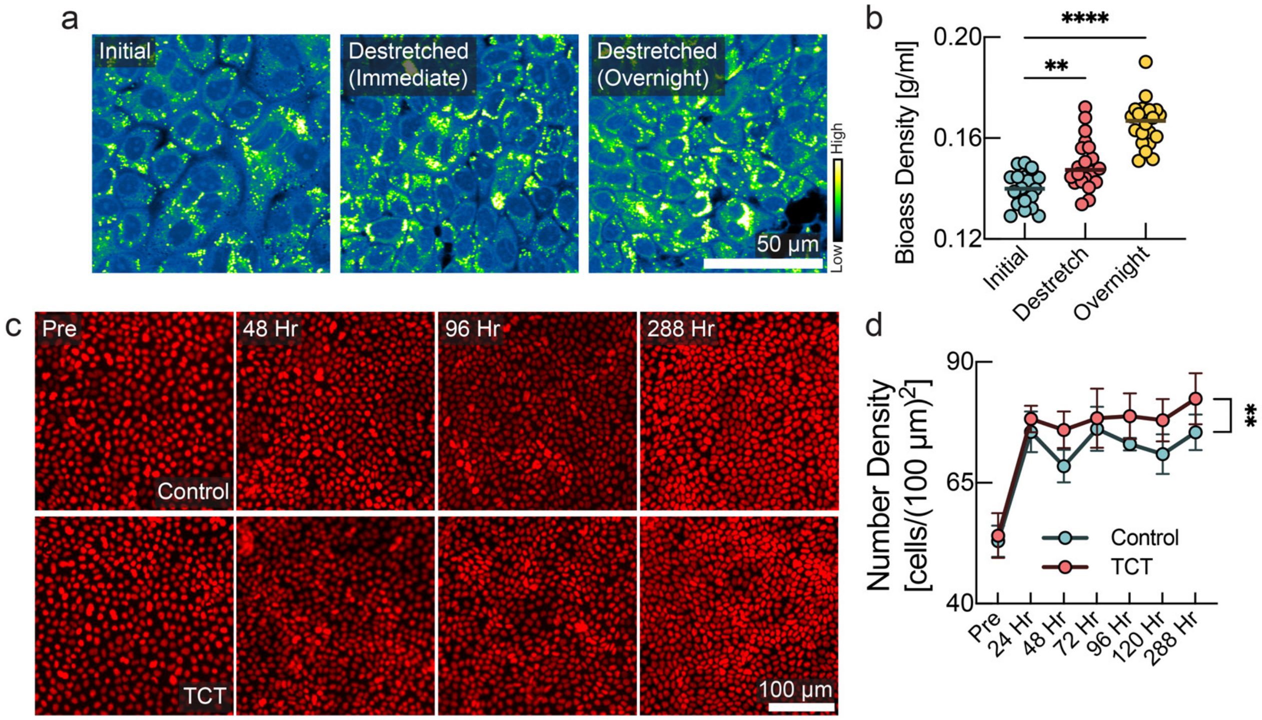


***Fig. S6: a-b, Modulation of cellular biomass density and cell-extrusion upon compression a.*** *Cells were grown to confluence on a pre-stretched stretchable membrane. Then the membrane was de-stretched uniaxially (20%) to introduce crowding. NoRI images representing the biomass density of initial tissue (left panel), immediately after compression (middle panel) , and overnight after compression (right panel) .* ***b,*** *Quantification of panel a show a significant increase in biomass density induced by mechanical crowding. (line at mean, n=20, p-values: 0.0013 (initial vs destretch), <0.0001 (initial vs overnight), unpaired t-test).* ***c,*** *Depolarization results in higher cell number density after destretching. MDCK cells were grown on stretchable membrane to confluence. Artificial crowding was induced via a 20% uniaxial de-stretch of the membrane. One set of membranes was kept under mild depolarizing conditions via addition of TCT, and another set was treated with the DMSO control. Nucleus images of both sets are shown.* ***d,*** *Quantification of number density for 12 days (mean ± s.d., n=10, p-value: 0.0032, unpaired t-test). De-stretching induced crowding in both cases, but cells reached a higher steady-state cell number density when treated with TCT.*

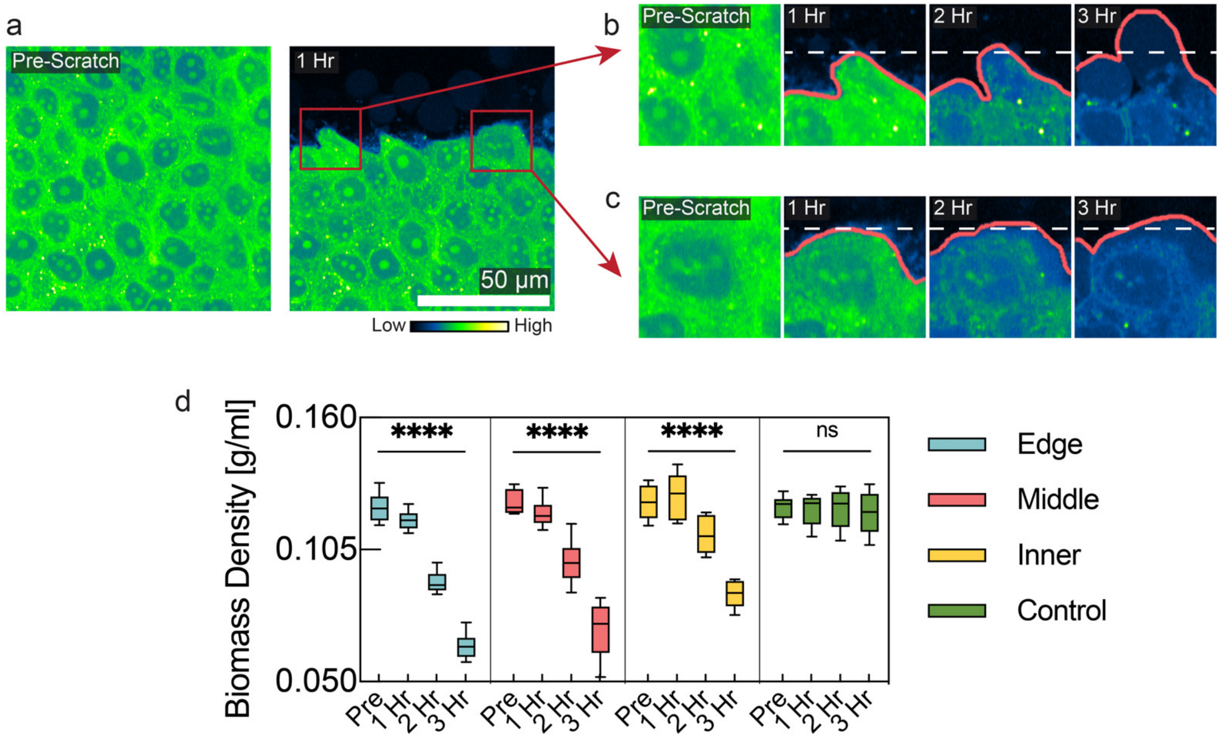


***Fig. S7: Cells expand at wound border with concomitant drop in biomass density. a,*** *NoRI images showing biomass density of MDCK monolayer before scratch and 1 hr after scratch.* ***b-c,*** *Inset kymographs images showing individual cells with expanding filopodia (indicating epithelial to mesenchymal transition) over time and subsequent dilution of intracellular biomass density. d, Quantification of cellular biomass density as a function of time and distance from scratch wound. Cells immediately next to scratch wound (cyan) exhibited an almost 50% reduction in biomass density over time. Cells further away from the scratch wound (inner and middle cells, yellow and red) exhibited a smaller and delayed drop in biomass density. The control area was chosen on the same plate, but at a distant location, centimeters away from scratch wound (box plot, line at mean, whiskers represent min to max, n=8).*

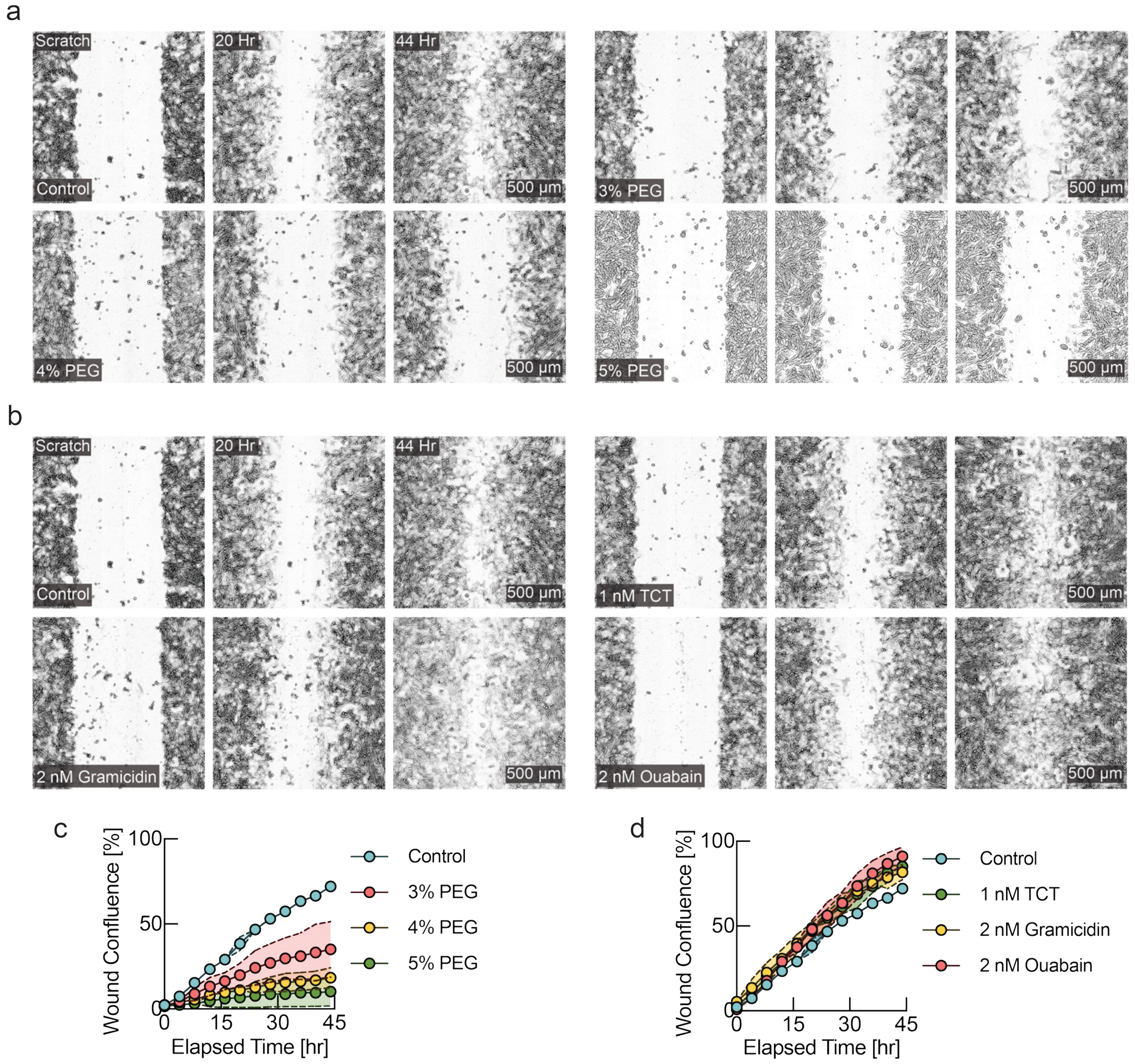


***Fig. S8: Wound healing efficiency is modulated by osmotic pressure and membrane potential.*** *CHO K1 cells grown on image lock plate with 96 wells. Uniform scratch wound was induced with an automatic scratch wound maker.* ***a,*** *Time-lapse images of the scratch wound assay with different concentrations of polyethylene glycol (PEG). Osmotic compression results in hyperpolarization (see Fig. S4) and negatively affected wound healing efficiency in a concentration-dependent manner.* ***b,*** *Time lapse images with various depolarizing drugs. Mild depolarization of membrane potential by various depolarizing drugs resulted in faster wound healing.* ***c-d,*** *Quantification of wound area closure as a function of time for panels a-b (mean ± s.d., n=2).*

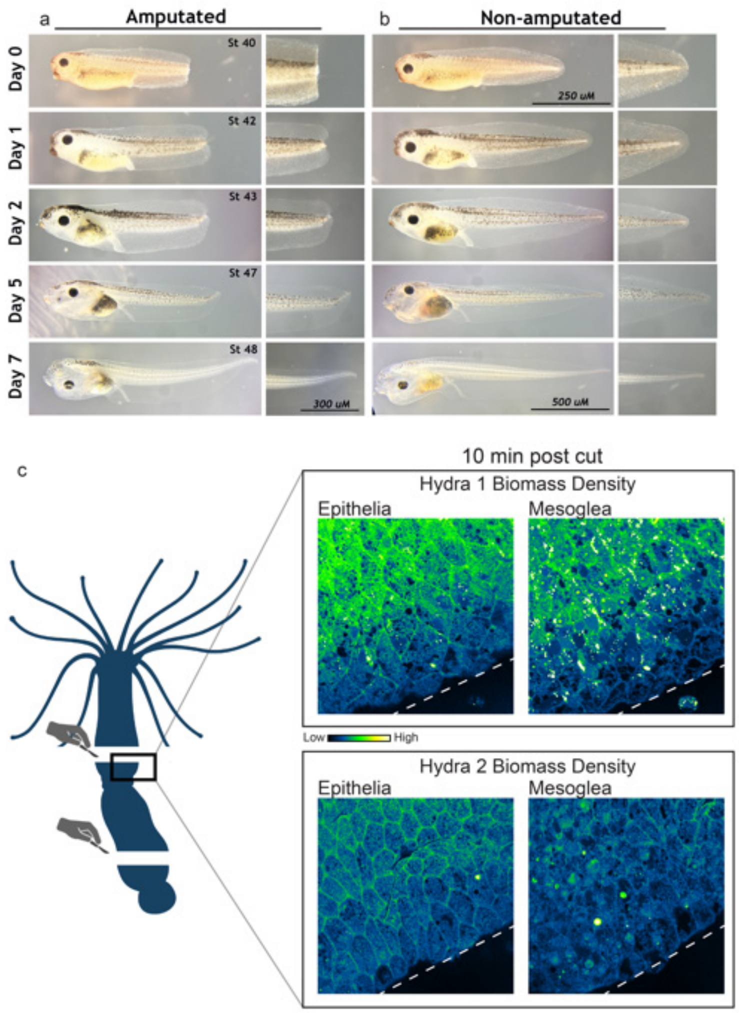


***Fig. S9: Tadpole tail and hydra regeneration assay***. *Stage 40 tadpole naturally regenerates tail after seven days post-amputation.* ***a****, Regeneration timecourse of an amputated tadpole and a highlight of the distal tail. Stage 40-amputate tadpole at the moment of tail cut (Day 0). Twenty-four hours later (Day 1), stage 42 tadpole displays a growth cone and remodeling of the superior and inferior distal fins. Stage 43 (Day 2, 48 hr after amputation) tadpole exhibits visible initiation of the tail regeneration. Stage 47 (Day 5) tadpole already displays a short forming tail. Seven days (Day 7) post-amputation, a regenerated tail is comparable to the sibling stage 47 tadpole. b, Non-amputated sibling tadpole and its highlighted distal tail from stage 40 to stage 47 grown in the same conditions as amputated tadpole.* ***c,*** *Gradient of biomass density is evident during hydra regeneration. Schematic shows experimental design. Cells at the wound edge were imaged using NoRI 10 minutes after amputation at both epithelial and mesogleal layers. Both epithelial and mesogleal cells showed a gradient of biomass density dilution across several cell layers away from the wound. White dotted lines represent the wound edge.*

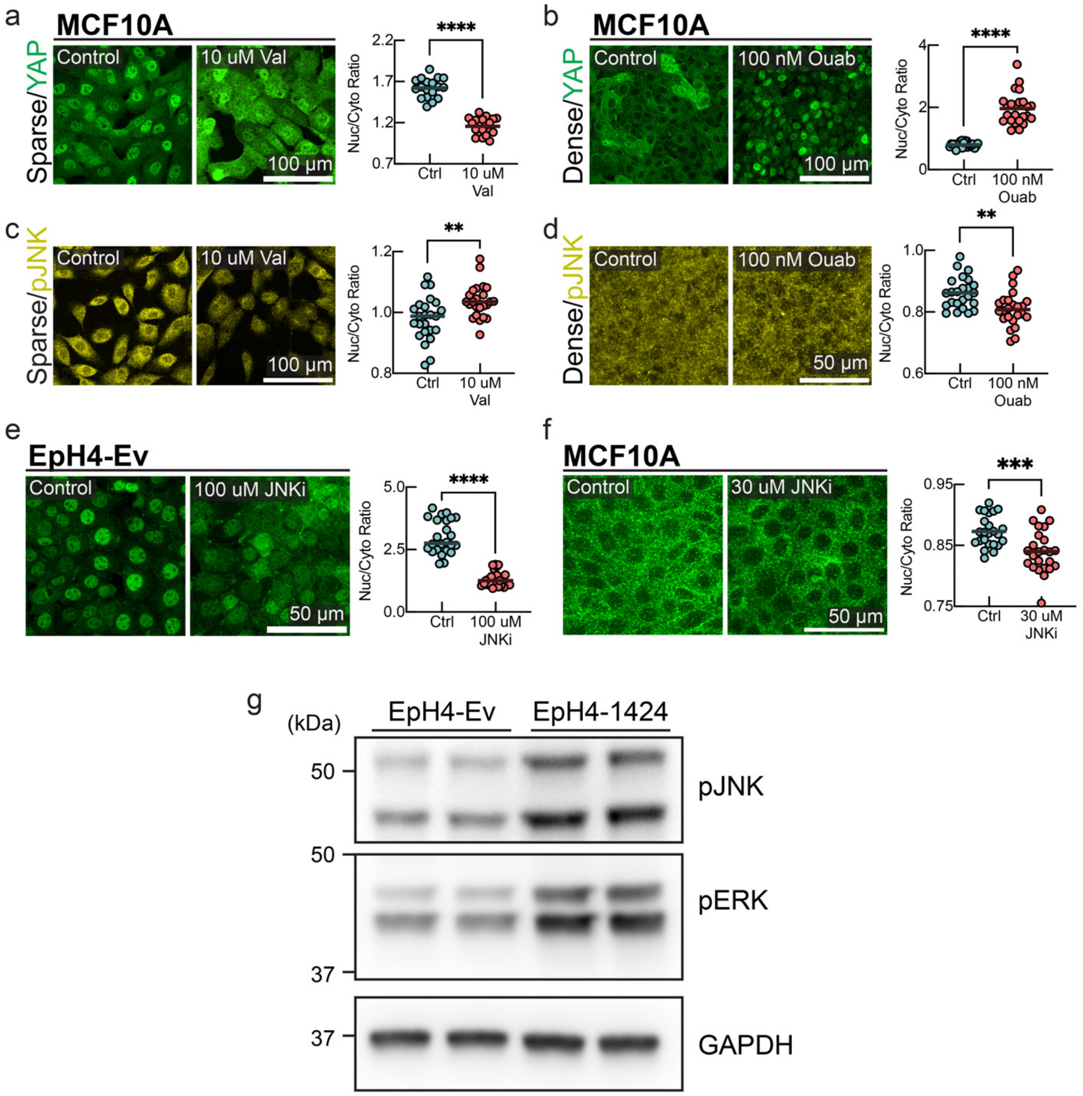


***Fig. S10: a-b**, YAP nuclear localization can be altered by changing the membrane potential in MCF10 cells. **a**, in sparse cells, induction of hyperpolarization by valinomycin causes significant nuclear exclusion as compared to the control. **b**, in dense confluent cells, depolarization of the membrane potential by ouabain prevents the nuclear exclusion of YAP. **c-d**, phosphorylated JNK nuclear to cytoplasmic ratio is modulated by membrane potential. **c**, pJNK predominantly localizes to cytoplasm in sparse cells, induction of hyperpolarization by valinomycin increases nuclear localization of pJNK in sparse cells. **d**, in confluent dense cells pJNK nuclear localization increases, induction of depolarization by ouabain promotes nuclear exclusion of pJNK in dense cells. **e**, Inhibition of JNK causes nuclear exclusion of YAP in EpH4. YAP immunostaining images of sparse EpH4-Ev cells under control and JNK inhibitor conditions, with quantification (line at mean, n=24, p-value <0.0001, unpaired t-test). **f**, Inhibition of JNK in dense confluent MCF10 cells, augments the nuclear exclusion of YAP. **g**, EpH4-1424 cells, which express constitutively active MEK (MEKDD) exhibit higher levels of pERK, and pJNK as compared to EpH4-Ev (expressing empty vector, no active form of MEK is expressed).*

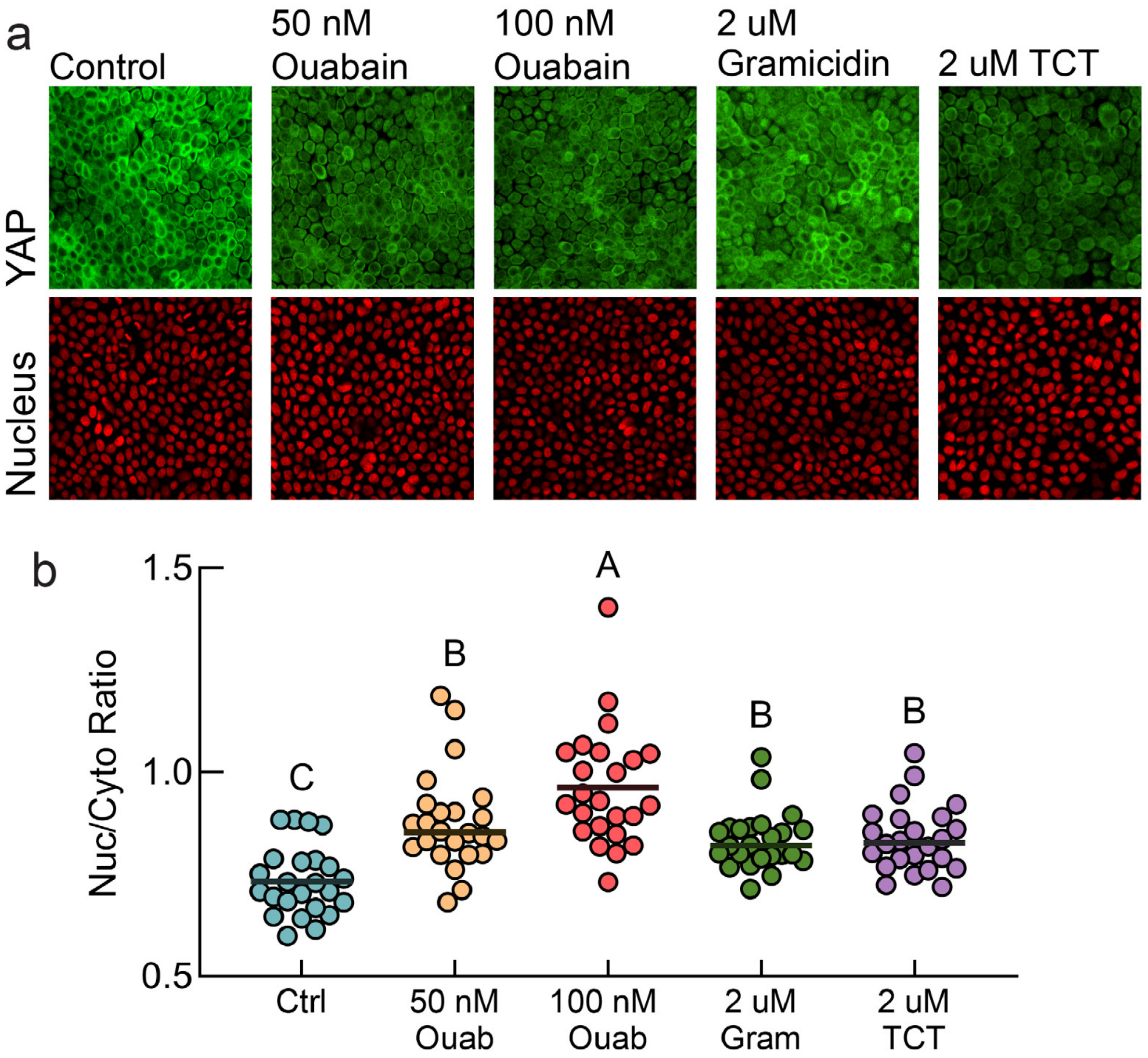


***Fig. S11: Depolarization causes YAP nuclear re-entry in dense cells. a,*** *YAP immunostaining images of dense MDCK cells under control and various depolarizing conditions* ***b,*** *Quantification of panel a (line at mean, n=24, compact letter display of multiple comparisons, Fisher's LSD test).*

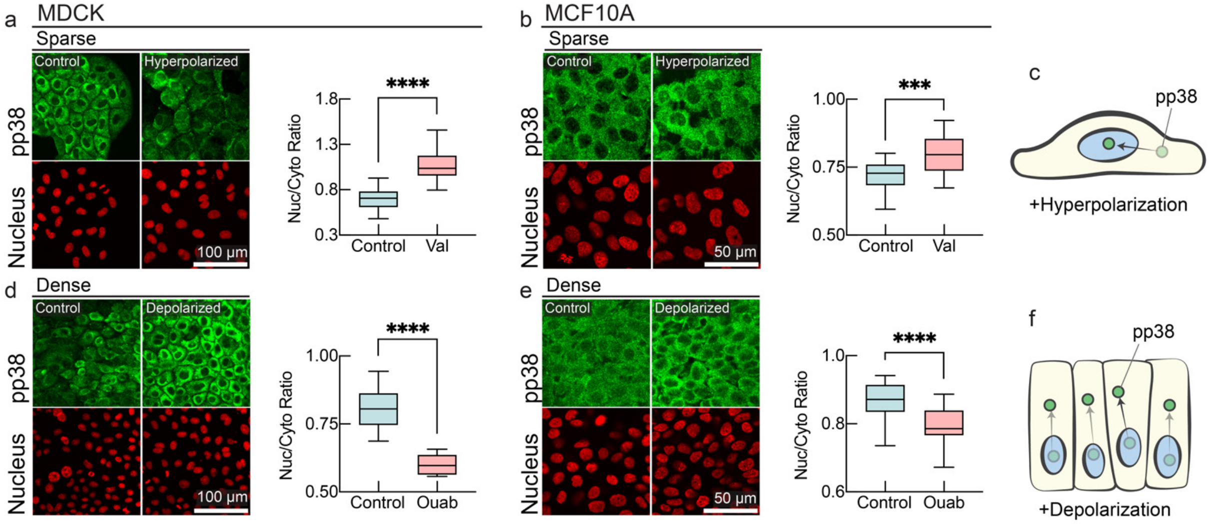


***Fig. S12: MAPK signaling is modulated by membrane potential.*** *a-f, Membrane potential dependence of localization of phosphorylated p38 (pp38) in immunostaining of MDCK and MCF10 cells. pp38 localizes predominantly in cytoplasm in sub-confluent (sparse) cells, and hyperpolarization by valinomycin results in augmented nuclear localization of pp38 in MDCK and MCF10 cells (panels a-b). pp38 localizes to cell nuclei in dense tissues, and depolarization by ouabain results in nuclear exclusion of pp38 (panels d-e). Nuclear to cytoplasmic ratio of pp38 is quantified in accompanying plots (box plot, line at mean, error bars represent min to max, n=24, p-values: <0.0001 (MDCK sparse control vs valinomycin), <0.0001 (MDCK dense control vs ouabain), 0.0001 (MCF10A sparse control vs valinomycin), <0.0001 (MCF10A dense control vs ouabain)).*

#### Materials and methods:

##### Cell Lines:

MDCK cells used in this study were obtained as a gift from Eugenia Piddini lab, current address, Bristol, UK. Another batch of MDCK cells were obtained from J.J. Fredberg lab, Harvard T.H. Chan School of Public Health. Canine origin of MDCK cells used in this study were verified by bulk RNA sequencing in the lab. MDCK cells expressing nuclear localized H2B:mCherry were made in lab by lentiviral transduction. CHO K1 cells were obtained from the HMS Systems Biology departmental freeze-down collection, which were originally obtained from ATCC. MCF10A H2B:mCherry cells were received as a gift from Caitlin Mills (Laboratory of Systems Pharmacology, Harvard medical school). Eph4-Ev cells were obtained from ATCC.

##### Culture media:

Cells were maintained in regular T25 and T75 tissue culture flasks (BD Falcon) for everyday maintenance. MDCK and EpH4-Ev cells were maintained in high glucose DMEM, supplemented with 10% bovine growth serum (Gibco), GlutaMAX (Gibco) and antibiotic-antimycotic solution (anti-anti, Gibco). CHO K1 cells were cultured in DMEM-HamF12 1:1 mixture with same supplements as described above. MCF10A cells were cultured in DMEM/F12 media supplemented with 5% horse serum (Gibco); recombinant human EGF (Proteintech, HZ-1326), final concentration 20ng/ml; Cholera toxin (Sigma-Aldrich, C8052-.5MG), final concentration 100ng/ml; Hydrocortisone (Millipore sigma, H0888-1G), final concentration 0.5 μg/ml; recombinant human insulin (Millipore-Sigma), final concentration 10μg/ml. GlutaMAX (Gibco) and antibiotic-antimycotic cocktail anti-anti (Gibco) was also added to the media.

##### Cell culture method:

For maintenance, cells were passaged regularly and subcultured in a timely manner. For experiments with fluorescence microscope, media was changed to FluoroBrite DMEM (Gibco), supplemented with 10% bovine growth serum, and antibiotic-antimycotic cocktail immediately before the imaging experiments.

##### NoRI imaging for biomass density measurement:

MDCK cells expressing nuclear localized H2B cherry were seeded at 0.25 x $10^6$ cells/plate on 60mm glass bottom plates. Cells were allowed to reach the onset of confluence for two days before imaging. We measured biomass density and cell number density from the same experiment using NoRI microscopy. Our custom-built NoRI imaging set up[9] also has fluorescence imaging abilities. Cell number density was calculated by quantifying number of nuclei per unit area, using 'Stardist' plugin[10] in Fiji. Biomass density of the cells was quantified from the NoRI images with a custom script, following the exact method described in detail in a previous work[9].

##### Biomass density measurement using holotomography:

To measure biomass density using holotomography, we used Tomocube microscope (Tomocube INC, South-Korea). This method allowed us to reconstruct the volume of the cell and measure total cellular biomass. MDCK H2B:mCherry cells were cultured on 60mm glass bottom plates and holotomographic images were taken. Cell nuclear images were also taken in the same experiment, using the fluorescence imaging mode of the Tomocube microscope. Cell number density was calculated by counting the number of cell nuclei per unit area with Stardist. For biomass density measurement, we segmented out a cubic volume inside the cell using Tomocube's image analysis software (Tomostudio). Total biomass of the cubic volume was calculated, and mass per unit volume was taken as the biomass density.

##### Altering tissue density by modulation of membrane potential:

For this experiment, MDCK H2B-mCherry cells were cultured on 96-well glass bottom plates. Cells were allowed to grow to homeostatic density, and then DMSO control, depolarizing and hyperpolarizing drug treatments were added to respective wells. We continued to change media everyday with respective added drugs to ensure drugs were not depleted or degraded. Plates were imaged using an Operetta High Content Imaging setup (Operetta, Perkin Elmer). Cell number per well was counted by segmenting and counting cell nuclei using the high content image segmentation pipeline available via the Columbus online server (Perkin Elmer).

##### Experiments on stretchable membrane:

PDMS based cell stretching chambers were procured from Strex-cell. We obtained two types of cell stretching chambers, either with 4 $cm^2$ surface area (Strex-cell STB-CH-04) for cell growth, or with 10 $cm^2$ surface area (Strex-cell STB-CH-10). NoRI experiments were done using the 10 $cm^2$ cell stretching chamber, and all other experiments were done using the 4 $cm^2$ cell stretching chambers. For all experiments stretching chambers were coated with 0.3 mg/ml type I bovine collagen (Advanced biomatrix, 5005), overnight at 4˚C. Membranes were washed 3 times with PBS before seeding cells.

To keep cell stretching chambers in stretched condition, we designed custom built holders, which were cut out of acrylic boards using a laser cutter from an HMS instrument facility. For short-timescale stretching and destretching experiments (Fig. 2b,e), we used a screw-based cell stretching system obtained from Strex-cell which (Strex-cell ST-0040).

Cell stretching chambers were first mounted on holders (either acrylic based or screw based), and cells were seeded on collagen coated cell-stretching chambers and kept in 150mm tissue culture dishes and placed in a humidified 37˚C incubator when not being imaged. Cells were allowed to grow to reach confluence with change of media every day.

##### Membrane potential measurement upon cell stretching and de-stretching:

For membrane potential measurements, we used the cell stretching chambers with 4$cm^2$ surface area and screw-based manual stretching device for continuous stretching. Cell stretching chambers were mounted on the stretching device in un-stretched condition and coated as outlined above. In one experiment, we wanted to determine how fast membrane potential responds on mechanical forces. To measure membrane potential, we used FluoVolt, a fast-responsive membrane potential sensitive dye (FluoVolt membrane potential kit, Thermo F10488). Before imaging, the media was changed to FluoroBrite DMEM (Gibco) containing FluoVolt membrane potential sensitive dye, following the protocol and reagents provided in the kit. Time-lapse microscopy images were recorded using a spinning disc confocal microscope. Images were taken before and after application of a 20% uniaxial stretch.

In a separate experiment, we wanted to check if membrane potential changes are reversible upon stretching and de-stretching. For this experiment we chose a different membrane potential sensitive dye $DiSBAC_2(3)$ (), which has high dynamic range and is suitable for detecting both depolarization and hyperpolarization in the same experiment. Before imaging, the media was changed to FluoroBrite DMEM (Gibco) and the dye was added to the media to achieve a final concentration of 5µM. Timelapse microscopic images were recorded using a spinning disc confocal microscope. Images were taken before stretching, after application of 20% uni-axial stretch and once again after de-stretch.

##### Preventing cell elimination on crowding by depolarization:

MDCK H2B:mCherry cells were seeded on collagen-coated 4 $cm^2$ chambers and allowed to reach early confluence with regular change of media. Cell nuclear images were taken using a spinning disc confocal microscope in stretched condition. After that, the stretched chamber was released from one end of the holder to induce crowding. Another set of images of the cell nuclei was taken immediately after destretching the membrane. In one batch of membranes, 100nM Ouabain (Sigma) was added at this stage. In the control group, DMSO was added to the media. Both groups of cells stretching chambers were transferred back to the tissue culture incubator and incubated overnight. Another batch of images was taken the next morning (~17 hours). Cell number density was quantified by counting number of nuclei per unit area using the Stardist Fiji plugin.

In a separate experiment, crowding was induced when cells were still in growth phase and a different depolarizing drug (TCT[11,12]) was added. In this experiment, cells growing on cell stretching chambers were maintained for 12 days and nuclear images were recorded intermittently, with regular media changes.

##### Membrane potential measurements in scratch wound:

MDCK cells were grown on 60mm glass bottom plates and allowed to reach early confluence. Before the experiment, media was changed to Fluobrite DMEM and 5µM $DiSBAC_2(3)$ membrane potential dye was added to the media. Images were taken using a spinning disc confocal microscope equipped with an on-stage incubator (Okolab, Japan). CellMask (Thermo) was added to media to stain the cell boundaries. Time-lapse images of membrane potential dye and cell plasma membrane were taken using separate laser illumination and filters. Scratch wound was created using a needle before time-lapse images were taken.

##### Biomass density measurement in scratch wound:

MDCK cells were seeded on 60mm glass bottom plates and NoRI images were recorded as mentioned before. Scratch wound was induced with a needle and time-lapse NoRI imaged were taken.

##### Immunostaining:

MDCK H2B:mCherry cells were seeded on 24 well glass bottom plate (MattTek) at 0.025 x $10^6$ cells/well and 0.05 x $10^6$ cells/well for dense conditions and waited for two days. Afterwards, cells were treated with depolarizing (ouabain, gramicidin) or hyperpolarizing (valinomycin) drugs or DMSO control. We changed the media in each well with the drug containing media or the DMSO control media, and cells were incubated with respective treatment overnight. The next morning, cells were fixed with 4% formaldehyde (diluted in 0.2% Tween-20 containing TBS (TBST)) for 20 minutes and washed for 5 minutes with TBST 3 times. The cells were then blocked in 1% BSA (diluted in TBST) for 60 min, washed again for 5 minutes with TBST for 3 times, and incubated in primary antibody diluted in 0.1% BSA-containing TBST. Primary antibody was removed the next morning, and 3 5-minute washes in TBST were given. Alexa-fluorophore coupled secondary antibodies were used for detection. Cells were imaged using a spinning disc confocal microscope.

##### Western blotting

MDCK cells were grown on plastic bottom 6 well plates, and treated with 5nM ouabain, 5nM gramicidin, 5nM valinomycin, or DMSO control for 48 hours. For total cell lysate preparation, cells were given 3 washes with ice cold PBS and lysed in RIPA lysis buffer (Millipore sigma), with protease and phosphatase inhibitor cocktail. Each lysate sample was passed 5-6 times through a 25G fine needle to shear DNA. Cell lysate was boiled with 4X LDS sample loading buffer (Invitrogen) and ran on pre-cast 4-12% bis-tris acrylamide gels (Nu-Page, Invitrogen). After electrophoresis, separated proteins were transferred on PVDF membrane using Invitrogen iBlot2 transfer apparatus and probed with respective antibodies. For detection, HRP-coupled anti-mouse and anti-rabbit secondary antibodies (Invitrogen) were used and blots were imaged in a Bio-Rad chemiluminescence imager using SuperSignal West Pico Plus (Thermo) chemiluminescence reagent.

##### Primary antibodies used:

Actin (C-2): sc-8432 (Santa Cruz Biotechnology). p-Src (Tyr 416) D49G4: #6943 (Cell Signaling Technology). YAP (63.7): sc-101199 (Santa Cruz Biotech). p-p38 MAPK (D-8): sc-7973 (Santa Cruz Biotechnology). p-JNK (Thr 183/Tyr 185) 81E11: #4668 (Cell Signaling Technology). MST1-2/STK3-4: A300-466A (Bethyl Laboratories Inc), FAT1: #10962 (Cell Signaling Technology), E-cadherin: 20874-1-AP (Proteintech), ZO-1 (R40.76): sc-33725 (Santa Cruz Biotechnology).

##### Secondary antibodies used:

Goat anti-mouse Alexa Fluor 405, Goat anti-rabbit Alexa Fluor 405, Goat anti-mouse Alexa Fluor 488, Goat anti-rabbit Alex Fluor 488, Goat anti-rabbit Alexa Fluor 647.

##### Drugs, inhibitors, and dyes used:

Ouabain octahydrate (Sigma-Aldrich, #O3125), Gramicidin A (Abcam, #AB144510), Valinomycin (Sigma-Aldrich, #94675). MST1 inhibitor XMU-MP-1 (Selleckchem, #S8334). JNK inhibitor, SP600125 (Selleckchem, #S1460). DiOC2(3) (3,3'-Diethyloxacarbocyanine Iodide, Thermo Scientific, #D14730). DiSBAC2(3) (Bis-(1,3-Diethylthiobarbituric Acid)Trimethine Oxonol, Thermo Scientific, #B413). DiSC3(5) (3,3'-Dipropylthiadicarbocyanine Iodide, Thermo Scientific, #D306).

##### FAT1 knockdown:

FAT1 SiRNA (Horizone/Dharmacon, ON-TARGETplus siRNA, #L-010513-00-0005) was transfected in MCF10a cells, following manufacturers protocol using the DharmaFECT 1 Transfection Reagent (Horizone/Dharmacon, #T-2001-01)

##### Xenopus tail regeneration:

Xenopus embryos were obtained by in vitro fertilization, cultured in 0.1 X MMR (NaCl 1 M, KCl 20 mM, $MgSO_4$ 10 mM, $CaCl_2 \cdot 2H2O$ 20 mM, HEPES50 mM, pH 7.4, supplemented with 1mM Kanamycin), and staged according to Nieuwkoop and Farber (1967)[13]. At stage 40, tadpoles were amputated, according to[14] Beck et al., 2012. Briefly, tadpoles were anesthetized in 0.1 x MMR containing 200 µM Tricaine for 1 min. The last third of the tail was amputated using a double-edged razor blade. Tadpoles were immediately transferred to a culture dish and regenerated for seven days at 18˚C. For NoRI image analysis, tadpoles were fixed after amputation in MEMFA (0.1 M MOPS (pH 7.4), 2 mM EGTA, and 1 mM $MgSO_4$, 3.7% Formaldehyde) at time 0, 10, 15, and 20 min postamputation. Imaging chamber on regular glass slide was created with Grace bio lab spacers (SecureSeal™ Imaging Spacers, Grace Bio Labs, SKU: 654002), and embryos placed within the imaging chamber after removing upper part of the body in 0.1x MMR. Imaging chamber closed with cover glass and imaged in NoRI microscope.

For depolarization analysis, embryos were amputated under anesthetics, and immediately transferred to 0.1X MMR with voltage sensitive dye (10 µM DiSBAC$_2$(3)). Embryos were allowed to recover for 10 minutes in 0.1X MMR and re-anesthetized with dye containing Tricaine. For imaging anesthetized embryos were placed in Grace bio labs imaging chambers with dye containing 0.1x MMR after removing upper part of the body. Imaging chamber was closed with a cover glass and imaged in an inverted spinning disc confocal microscope.

##### Hydra:

Wildtype Hydra was obtained from Florian Engert lab (Harvard) and maintained in packaged spring water following standard protocol. For regeneration assay and Raman microscopy, body trunk was amputated with a scalpel under dissection microscope. For imaging, a imaging chamber was made on regular glass slide using grace bio lab spacers (SecureSeal™ Imaging Spacers, Grace Bio Labs, SKU: 654002). Amputated body part was placed inside the imaging chamber in 10 µL water and a cover glass was placed to close the chamber. The slide was then imaged in the NoRI microscope.

#### Supplementary References


1. Hille, B. Ion channels of excitable membranes. in (2001).
2. Chang, R. *Physical Chemistry for the Biosciences*. (2005).
3. Milo, R., Jorgensen, P., Moran, U., Weber, G. & Springer, M. BioNumbers—the database of key numbers in molecular and cell biology. *Nucleic Acids Res* **38**, D750 (2010).
4. Milo, R. & Phillips, R. Cell Biology by the Numbers. *Garland Science* (2015).
5. Dyer, K. F. The Quiet Revolution: A New Synthesis of Biological Knowledge. *J Biol Educ* **5**, 15–24 (1971).
6. Bi, D., Yang, X., Marchetti, M. C. & Manning, M. L. Motility-driven glass and jamming transitions in biological tissues. *Phys Rev X* **6**, (2016).
7. Zatulovskiy, E., Zhang, S., Berenson, D. F., Topacio, B. R. & Skotheim, J. M. Cell growth dilutes the cell cycle inhibitor Rb to trigger cell division. *Science (1979)* **369**, 466–471 (2020).
8. Cone, C. D. & Tongier, M. Contact inhibition of division: involvement of the electrical transmembrane potential. *J Cell Physiol* **82**, 373–86 (1973).

9. Oh, S. *et al.* Protein and lipid mass concentration measurement in tissues by stimulated Raman scattering microscopy. *Proceedings of the National Academy of Sciences* **119**, (2022).
10. Schmidt, U., Weigert, M., Broaddus, C. & Myers, G. Cell Detection with Star-Convex Polygons. in 265–273 (2018). doi:10.1007/978-3-030-00934-2_30.
11. Barboiu, M. *et al.* Polarized Water Wires under Confinement in Chiral Channels. *J Phys Chem B* **119**, 8707–8717 (2015).
12. Barboiu, M. *et al.* An artificial primitive mimic of the Gramicidin-A channel. *Nat Commun* **5**, 4142 (2014).
13. Gerhart, J. & Kirschner, M. *Normal Table of Xenopus Laevis (Daudin)*. (Garland Science, 2020). doi:10.1201/9781003064565.
14. Beck, C. W. Studying Regeneration in Xenopus. in 525–539 (2012). doi:10.1007/978-1-61779-992-1_30.
